# Varroa mites escape the evolutionary trap of haplodiploidy

**DOI:** 10.1101/2024.05.10.593493

**Authors:** Nurit Eliash, Tatsuya Endo, Xia Hua, Maeva A. Techer, Juliana Rangel, Huoqing Zheng, Evan P. Economo, Alexander S. Mikheyev

## Abstract

Genetic diversity is essential for populations adapting to environmental-changes. Following genetic bottlenecks, invasive species often have reduced diversity, yet must rapidly adapt to novel environments. This paradox is especially pronounced in parasites, which face repeated transmission bottlenecks and anthropogenic countermeasures. The challenge of maintaining diversity in the short-term intensifies in haplodiploid species, where males inherit and transmit only half of their mother’s genome, limiting effective population size. To examine how such species may maintain adaptability, we studied inheritance in Varroa mites (*Varroa destructor*), globally invasive honey bee parasite. Using three-generation pedigrees, we found that Varroa is not haplodiploid, as previously thought. Rather, females produce diploid sons who transmit either maternal allele to daughters. This system slows reduced heterozygosity loss under sib-mating, compared to ancestral haplodiploidy, increasing effective population size and adaptability. Our findings reveal a rare reversion from haplodiploidy, long considered an evolutionary end state, and suggest reproductive plasticity underlies the resilience of invasive parasites like Varroa.

## Introduction

Genetic diversity is the basis for biodiversity, being required for species to evolve in response to environmental changes^1^. For a given selection coefficient, the effectiveness of natural selection *vs.* genetic drift is determined by the effective population size^2^. While selection dominates at larger population sizes, beneficial alleles can be lost due to drift in small populations. Biological invasions are often accompanied by genetic bottlenecks that reduce effective population size. Parasites are particularly at risk, as they often experience bottlenecks as part of their life cycle: an introduction to a new host often involves one or a few individuals, and the host-parasite population dynamics also involve frequent and drastic changes in the population’s size^3^. In addition, because of difficulties involved in finding unrelated mates within a host, many parasites frequently inbreed, which further decreases genetic variability^4–6^. Although all the above scenarios are expected to decrease the effective population size, by definition, invasive parasites are successful species that adapt to different environments, hosts, and in many cases, the chemical pesticides used to control them^7,8^. This ‘genetic paradox of invasion’^9,10^ raises the question of how invasive species with expected low genetic diversity are so successful. A few solutions have been suggested for this paradox. These include physiological explanations^11–13^, and multiple invasions that compensate for the initial low genetic diversity^13–15^. Occasionally, organisms evolve unusual reproductive strategies that help them avoid inbreeding. For example, in some ants, males and females constitute genetically distinct clonal lineages that combine to produce sterile workers but otherwise do not intermix^16,17^. Detecting unusual reproduction mechanisms typically requires multi-generation-controlled crosses, which are often difficult if not impossible to recreate (especially in non-domesticated organisms), suggesting that many unusual reproductive modes remain to be discovered.

Here we examined the reproductive biology of the Varroa mite (*Varroa destructor*), a global invasive parasite of honey bees. These mites have experienced a range of genetic bottlenecks. They went through a genetic bottleneck when switching from their original host, the Eastern honey bee (*Apis cerana*) to the Western honey bee (*Apis mellifera*)^18^. They also lose genetic diversity when colonizing new hives. Lastly, they routinely sib-mate in small families (typically one male mates with three to four female sisters)^19,20^. This mating behavior further increases the strength of genetic drift, and makes natural selection less effective. Yet, Varroa shows high adaptability: it efficiently switches between host species^21–23^, and can rapidly evolve resistance to pesticides via various mechanisms^24^. While a high level of adaptability explains the mite’s successful global spread, the mechanisms underlying this adaptability remain puzzling in the light of regular genetic bottlenecks and routine inbreeding.

Varroa is believed to be haplodiploid, whereby fertilized eggs develop into diploid females, while unfertilized eggs develop into haploid males^20,25–27^. Haplodiploid species have a smaller effective population size relative to diplodiploid systems^28,29^, though haplodiploidy may confer long-term advantages in inbreeding systems such as more effective purging of deleterious alleles^30^. In Varroa, haplodiploidy was inferred from two karyotyping studies^31,32^. Yet, as those studies examined Varroa embryos only, sex could not be directly determined. Häußermann et al.^20^ found that unmated females can produce males— consistent with arrhenotokous parthenogenesis *vs.* paternal genome elimination (PGE) that is found in many other mites. Yet, while consistent with haplodiploidy, these lines of evidence are indirect.

Haplodiploidy is believed to be evolutionarily stable, sometimes being referred to as an ‘evolutionary trap’, suggesting limited potential for an evolutionary reversion to diploidy^33,34^. Cruickshank and Thomas^35^ used phylogenetic reconstruction to infer that Varroa evolved from diploid ancestors, via intermediate pseudo-arrhenotokous species. While Varroa was included in that study, its haplodiploidy was assumed based on the fact that all members of its family were haplodiploid. However, the taxon sampling was sparse and until recently, the phylogenetic placement of Varroa was uncertain, opening conclusions to criticism^36^. Indeed, more complete investigations were hampered by the uncertain taxonomic placement of Varroa, until a recent study placed it as a subfamily within Laelapidae, allowing a more targeted comparison^37^. Of the 19 sexually reproducing Laelapid mites in the Tree of Sex database^38^, only a single one is not haplodiploid: *Dicrocheles phalaenodectes*, a parasite inhabiting the ear canal of some Noctuid moths. However, the placement of *Dicrocheles* in Laelapidae is based solely on morphological data, and has been debated^39^; it is certainly only distantly related to Varroa. Thus, based on the sum total of the evidence, haplodiploidy was ancestral to Varroa.

Theory suggests that reversal to diploidy from arrhenotoky is unlikely^34^, and, indeed, no reversals of arrhenotokous sex determination are known to date^40^. Yet, despite strong circumstantial evidence suggesting the existence of arrhenotokous parthenogenesis in Varroa, no study has examined Varroa’s mode of genetic inheritance, making its success in spite of genetic handicaps somewhat of a mystery. Here, we explicitly examine this species’ mode of genetic inheritance using multi-generational pedigrees.

## Results

Two lines of evidence suggest that Varroa males are functionally diploid: [1] their somatic cells inherited and expressed both maternal alleles (Fig 1 and 2), and [2] they randomly transmited these alleles to their daughters (Fig 1). Modeling shows that this increases the mite’s effective population size relative to ancestral haplodiploidy (Fig 3).

**Figure 1.**
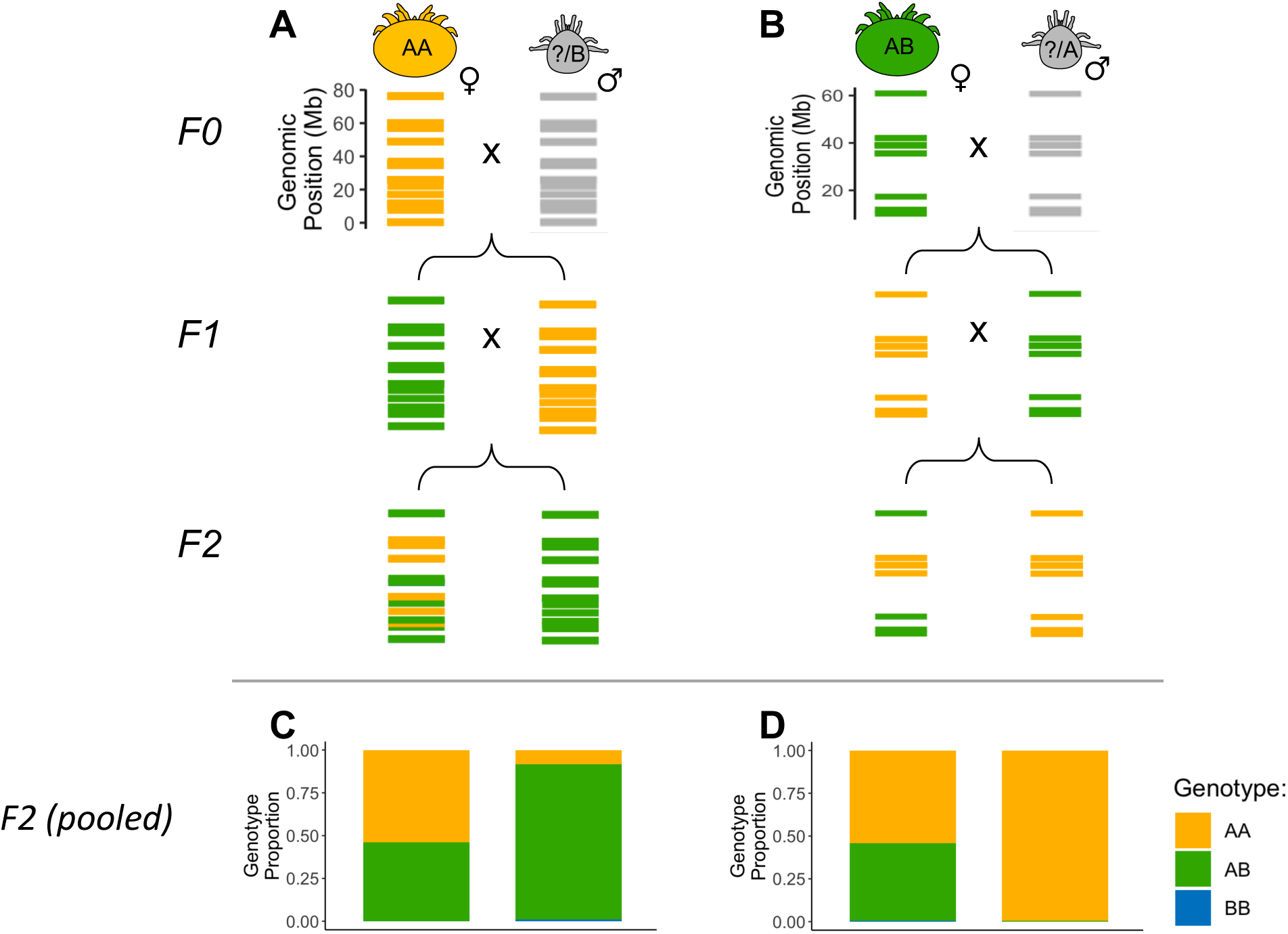
**Inheritance patterns in *Varroa destructor* mites shows that males inherit two alleles from their mother and transmit both of them to their daughters with equal probability. Females are produced sexually. Panels A and B show the genotype flow of sites on the largest chromosome, NW_019211454, for a representative family spanning three generations: the foundress grandmother (F0 female), her offspring (F1 female and male), and their F2 offspring (F2 female and male).** [Panel A]: Family 478, crossing a heterozygous F1 female (AB) with a homozygous F1 male (AA), showing the genotypes of their F2 female and male offspring across 33 sites on chromosome NW_019211454. [Panel B]: Family 534, crossing a homozygous F1 female (AA) with a heterozygous F1 male (AB), showing the genotypes of their F2 female and male offspring across 14 sites on chromosome NW_019211454. **Panels C and D show the pooled genotype proportions in F2 females and males from all families, across all sites on all seven chromosomes, for the same cross types as in A and B, respectively.** [Panel C]: Pooled data for the cross of heterozygous F1 female (AB) × homozygous F1 male (AA): 7 F2 females (11,207 sites) and 12 F2 males (16,515 sites). [Panel D]: Pooled data for the cross of homozygous F1 female (AA) × heterozygous F1 male (AB): 11 F2 females (4,999 sites) and 10 F2 males (2,536 sites). F2 females show a ∼1:1 ratio of AA:AB genotypes, close to sexual reproduction, while F2 males predominantly inherit their mother’s genotype, suggesting maternal inheritance. The data are filtered using strict quality thresholds (VAF < 0.01 for AA, 0.2-0.8 for AB, > 0.9 for BB; corresponding QAF thresholds). As expected from AA × AB crosses, no BB genotypes were observed. F0 males were not collected for the pedigree, as they remained in the cell when the honey bee emerges. However, based on the genotypes of the F0 and F1 offspring, we could infer the F0 male genotype. For instance, in the cross of F1 AB female × AA male, the F0 male must have had at least one B allele (indicated as “?/B” in panel A). In the cross of F1 AA female × AB male, the F0 male must have had at least one A allele (indicated as “?/A” in panel B). Colors: Orange = AA, Green = AB, Blue = BB. Note: “?” indicates an inferred, but unobserved allele.

**Figure 2.**
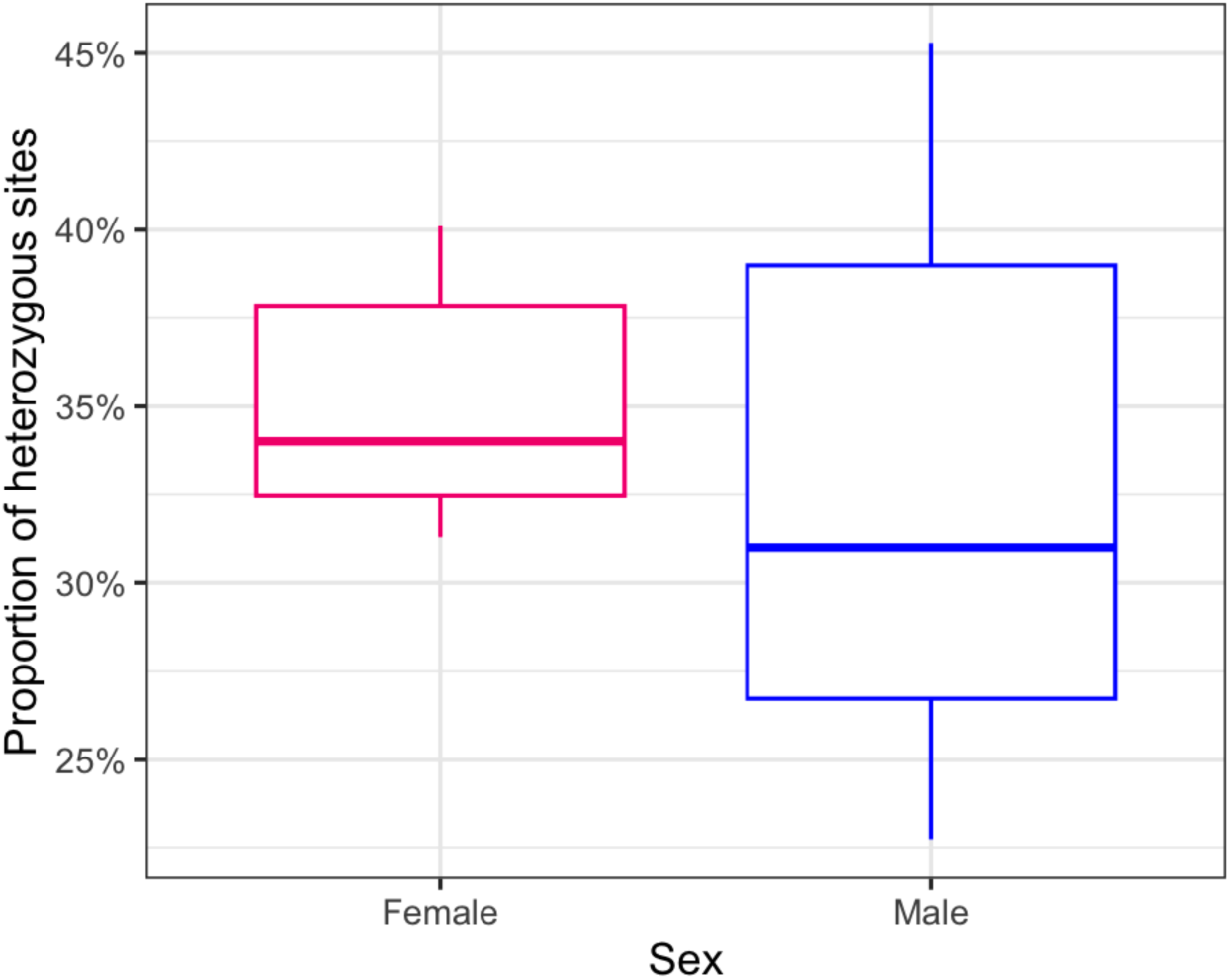
RNA-seq analysis of heterozygous site proportions in male and female Varroa mites. Each point represents the mean proportion of heterozygous sites per mite, with error bars showing 95% confidence intervals, and lines connecting mother-son pairs. The two sexes show similar heterozygosity proportions in their RNA across five families (mean difference = 0.022 ± 0.10, paired t-test: *t* = 0.46, df = 4, *P*-value = 0.672). The y-axis shows the proportion of heterozygous sites as a percentage of total sites analyzed. Males are much smaller than females, and result in lower-quality sequencing libraries, possibly explaining their greater variability. Note that, while males and females are clones, we do not expect identical heterozygosities in transcriptomic data because different genes are expressed in the two sexes.

**Figure 3.**
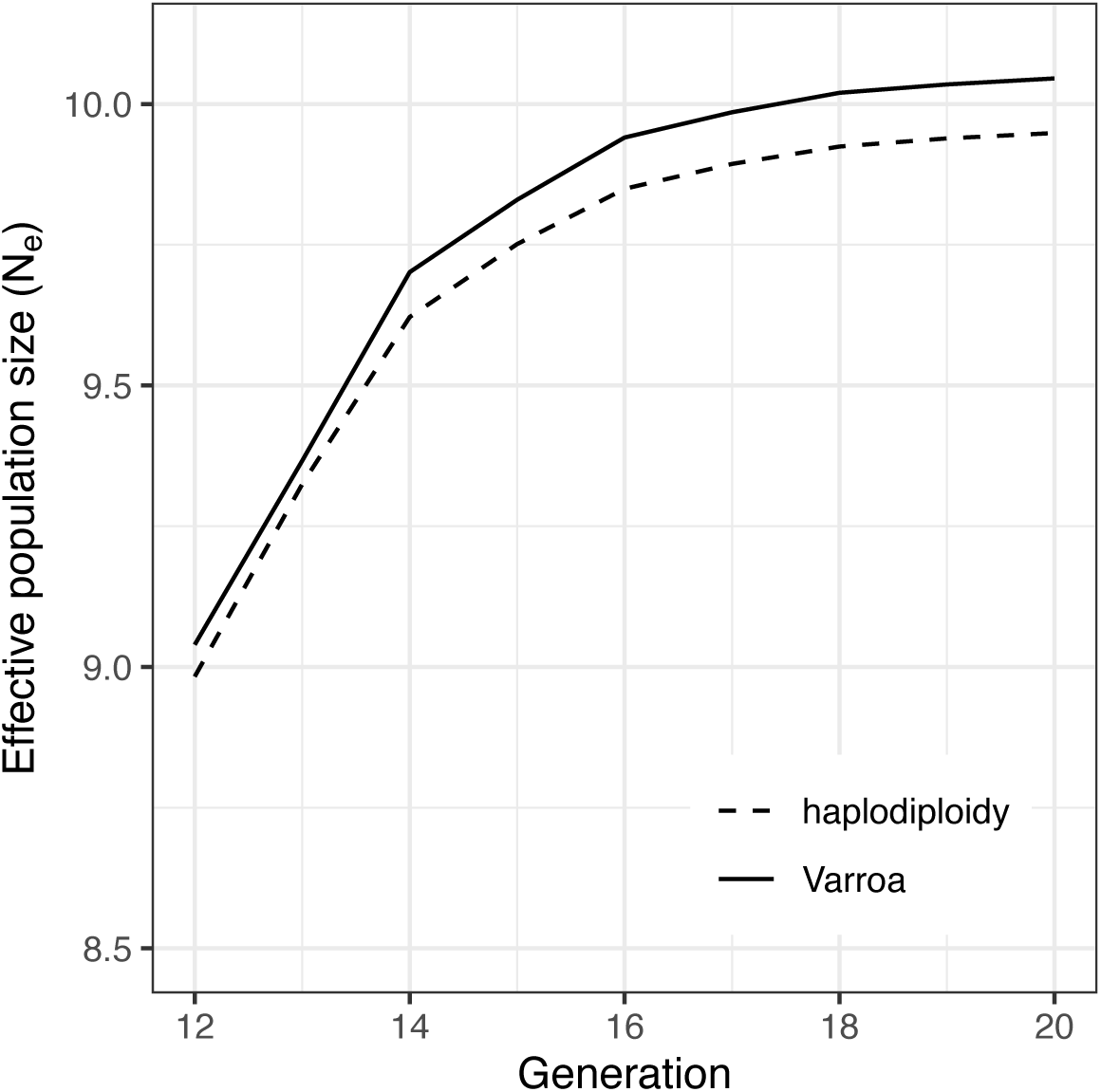
Varroa diploid arrhenotoky results in a greater population size, compared to ancestral haplodiploidy under identical population conditions. This is a representative plot simulated under the following conditions: 2 daughters per mother, a carrying capacity of 100 mites, and 0.5 initial correlation between gametes. The reduced correlation due to male diploidy produces a small but consistent increase in effective population size under inbreeding conditions.

### Males are produced by diploid arrhenotoky preserving their mothers’ genotype

Adult males inherited both of their mother’s alleles, as evident from the three-generational pedigree study. Looking at the cross between a heterozygous F1 female (AB) and a homozygous F1 male (AA), we found that all variants of the F2 grandson in a family maintained their mother’s heterozygosity (Fig 1A, example of family 478). This was consistent across families, as pooled data showed 97% of the genotypes in F2 males maintained their mother’s heterozygosity (Fig 1C). Similarly, when crossing a homozygous F1 female (AA) and a heterozygous F1 male (AB), variants of F2 grandson were homozygous like the F1 mother (Fig 1B), and the pooled-family’s data were in consistent (Fig 1D). Both alleles were also expressed, as evidenced from heterozygosity in RNA-seq data of adult female mites and their sons (Fig 2, Table S1). The two sexes showed similar heterozygosity proportions in their RNA across five families (mean difference = 0.022 ± 0.10, paired t-test: *t* = 0.46, df = 4, *P*-value = 0.672).

### Males transmit their two alleles to their daughters with equal probability

Males not only inherited two alleles, but they also randomly transmitted each allele to their daughters, in a typical Mendelian manner. Looking at the cross between a homozygous F1 female (AA) and a heterozygous F1 male (AB), we observed that about ∼50% of the variants in F2 females were homozygous (like mother) and the rest were heterozygous (like the father) (Fig 1B). F2 male offspring of the same cross were homozygous (AA), just like their mother. This was consistent across families, as shown by the pooled data (Fig 1D).

### All else being equal, Varroa diploid male arrhenotoky has greater population size relative to ancestral haplodiploidy

We conducted genetic transmission path modelling, which traced gene flow from ancestors to descendants and quantified genetic relatedness through path coefficients, following Wright ^41^. This showed that Varroa lost slightly but consistently less heterozygosity under inbreeding conditions, compared to haplodiploids, and therefore had a correspondingly higher effective population size (Fig 3). The difference arose because the two alleles in male genomes lowered the correlation between uniting gametes and attenuated inbreeding compared with haplodiploid mating, whereby the male inherited a single allele. Consequently, for the same census size, the limit of Ne was higher under diploid arrhenotoky across the explored ranges of mothers per generation, daughters per mother, and brood-cell occupancy probability.

## Discussion

In this study, we found that *V. destructor* has evolved from ancestral haplodiploidy ^35^ to a novel reproductive strategy with diploid arrhenotokous males that are clones of their mothers (see Fig 4 for a scheme of the inferred mode of genetic inheritance). This strategy potentially has both short-term, and long-term benefits for Varroa, each overcoming a different evolutionary ‘paradox’. In the short-term, the production of clonal males increases the females’ fitness by assuring transmission of both of her genetic variants. This strategy can overcome the “paradox of sex” by blending the benefits of sexual and asexual reproduction. Namely, we found that females can produce sons containing both copies of their genes, but female eggs are produced meiotically. In the long term, functional diplodiploidy can potentially increase the effective population size (vs. haplodiploidy) (Fig 3). A higher effective population size imparts greater evolutionary potential, and indeed Varroa has been remarkably adaptable despite repeated bottlenecks exemplifying the “genetic paradox of invasion”.

**Figure 4.**
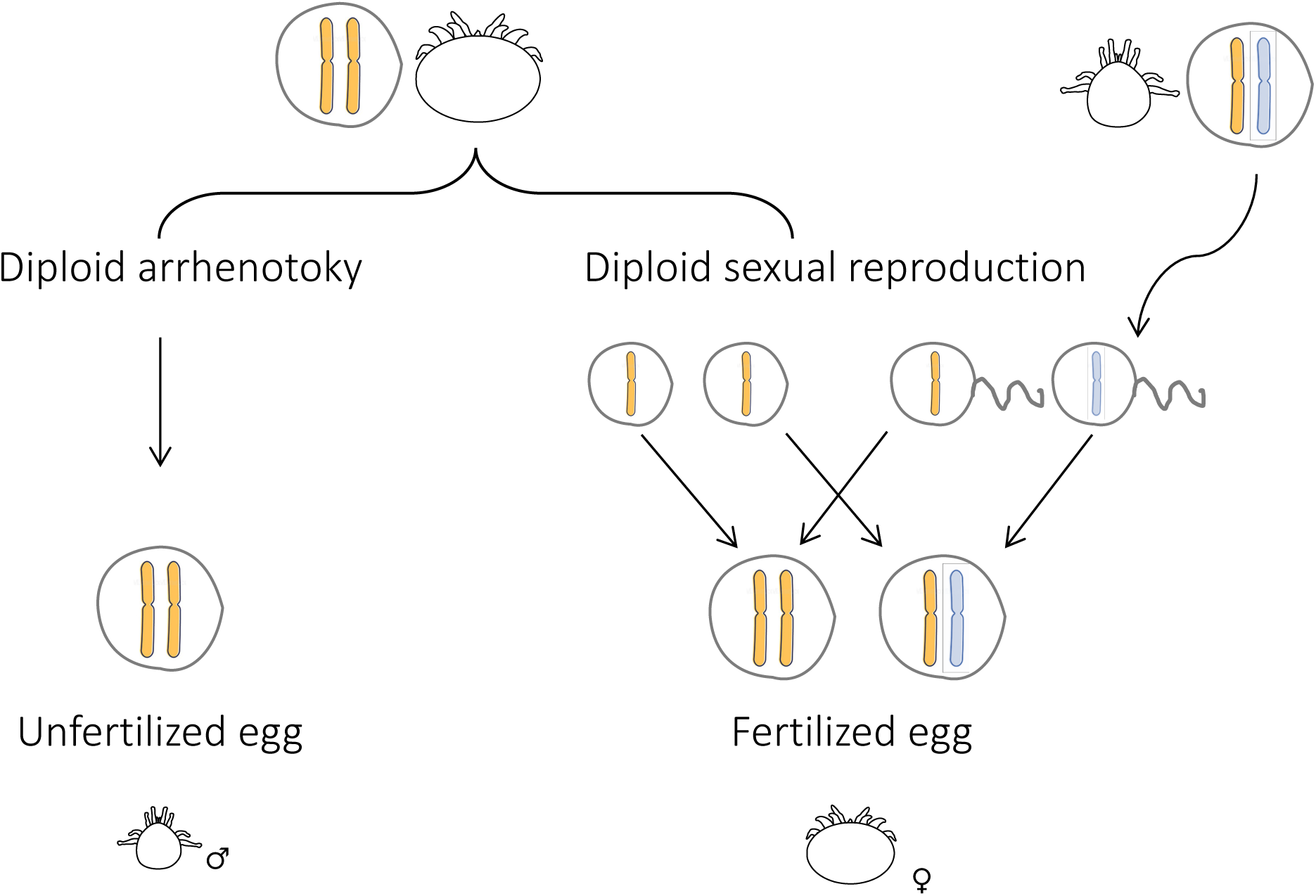
Hypothesized mode of genetic inheritance in Varroa male and female mites. Females are produced via diploid sexual reproduction, while males are produced via diploid arrhenotoky. To illustrate the inheritance of maternal and paternal alleles in the two sexes, we show an example of a cross between a homozygous female and a heterozygous male, and their possible female and male offspring (as in Figure 1).

Another evolutionary surprise from this study is the effective loss of haploid arrhenotoky. Once evolved, asymmetric inheritance appears to reach an evolutionarily stable state, sometimes referred to as an ‘evolutionary trap’^34^. Obstacles to re-evolving Mendelian inheritance are diverse. One obstacle is mechanistic, in that males would need to re-evolve meiosis, which was always thought to be difficult^42^. However, recent work from asexual stick insects indicates that male meiotic regulatory mechanisms are preserved in females^43^, so re-evolving male meiosis may be easier than expected. The other obstacle is evolutionary, as examined in detail by Bull^44^. In that scenario, females reach a fitness optimum in haplodiploid systems, being able to control male production and thus the sex ratio. Thus, re-emergence of diploid males would have to be a male trait, and biparental diploid males must carry mutations that allow them to produce diploid sons. However, given the level of arrhenotokous female control over fertilization and sex ratios, those traits are theoretically unlikely to spread^44^. Indeed, the only evolutionary feasible transition from haplodiploidy appears to be towards parthenogenesis, though these are short lived^45^. Yet, Varroa overcame the constraint on the production of diploid sons by having females solely control both, the sex ratio, and diploid male production. The production of meiotic sperm and the evolution of diploid arrhenotokous male production illustrate how the unforeseen complexity of biological mechanisms may challenge long-standing elegant mathematical models.

Although the mechanism through which sex is determined in Varroa still needs further investigation, other facultatively parthenogenetic systems provide insights into how it may hypothetically occur. For instance, Cape honey bees (*Apis mellifera capensis*) store new eggs in their oviducts arrested in metaphase I, while meiosis II takes place only after oviposition^46^. If the egg is not fertilized, it develops into a female by central fusion. If it is fertilized, a diploid female is sexually produced. A similar or parallel mechanism may be at play in Varroa, except that unfertilized eggs undergo central fusion to develop into males while preserving their mother’s genotype (Fig 1). Unlike honey bees, where sex is controlled genetically, an epigenetic factor would be required for sex determination in Varroa. Such a factor may be provided by the female, for instance by controlling fertilization.

### Conclusions

Our work underscores the importance of comprehensive experimental inquiries into diverse reproductive mechanisms, even in organisms that appeared to be well studied. Recent investigations have unveiled a broader spectrum of reproductive strategies than were previously recognized^42^, exemplifying the dynamic nature of biological systems. Yet, the discovery of a novel reproductive system, such as that in Varroa, can have wide-ranging fundamental and applied consequences. For instance, Varroa’s haplodiploidy is assumed when considering how it evolves in response to countermeasures, such as pesticides, breeding honey bees for parasite tolerance, or meiotic drive^47–49^. Furthermore, Varroa has emerged as a compelling model for the study of the evolutionary dynamics in reproductive systems, offering insights into mechanisms for overcoming evolutionary constraints and the intricacies of sex determination. Our findings open promising avenues for future research, contributing to a deeper understanding of evolutionary innovation and reproductive plasticity in Varroa and beyond.

## Materials and methods

### Honey bees and mites

#### For the pedigree study

Varroa mites and honey bees (*Apis mellifera* L.) were obtained from colonies at the Onna village apiary and the experimental apiary of the Okinawa Institute of Science and Technology (OIST) in Okinawa, Japan. All colonies used in the experiment were maintained without any treatment against Varroa and received a 60% w:v sugar solution and 70% pollen patties as supplemental feed when necessary. Experiments were conducted between July and November 2020.

#### For comparative transcriptomic analysis of Varroa gene expression heterozygosity in mothers and sons

Varroa mites were collected from *A. mellifera* colonies at the Zhejiang University in CITI, China, in November 2024. A total of ten mites were collected from five families, each consisting of a mother and her son.

### Artificial infestation for Varroa pedigree construction

To obtain a three-generational mite pedigree (F0, F1, and F2), we used a semi-natural infestation system. This started with the collection of a naturally infesting mated Varroa female (F0). Her adult daughter (F1) was then artificially introduced into a honey bee pre-pupal cell and allowed to commence the reproductive phase of its life cycle. The F1 mite and her offspring (F2) were then collected at the end of the reproductive cycle. For a detailed description of the experimental design, see Fig 5 and the sections below. Using this approach, we could genotype all members of the pedigree, except for the male that mated with the F0 female.

**Figure 5.**
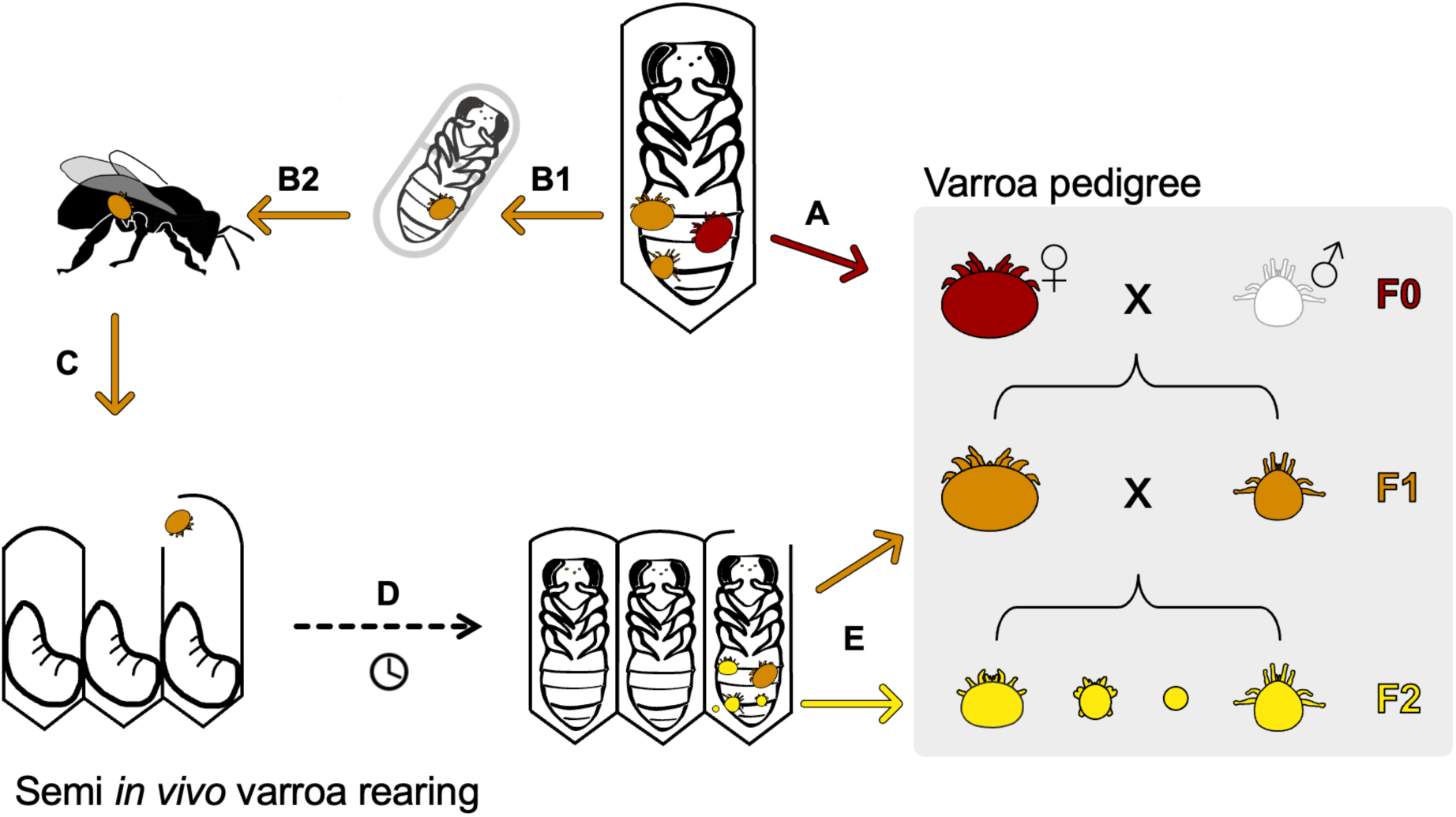
Pedigree experimental design. A three-generation Varroa pedigree (F0-in red, F1-in orange, and F2-in yellow) of 30 families was constructed via the following steps: **A.** Collecting a naturally infested family: foundress female mite (F0), and her two adult offspring: male and female F1 mites (F0 male’s genotype was not possible to study because the F0 female was previously mated, typically by a single male); **B.** “Incubation phase”: (B1) Transferring the young F1 female onto a bee pupa for 1-4 days, till cuticle sclerotized and darkened; “Dispersal phase”: (B2) Transferring F1 female onto a nurse adult bee for another 3 days; **C.** Introducing the matured F1 female to a freshly capped 5th instar larva cell; **D.** F1 female reproducing inside the bee cell; **E.** 10 days post introduction, collecting the artificially infested family: F1 female and her F2 offspring. The F2 generation included a son and one or more daughter nymphs. This design provided known parents and genotypes for the F1 and F2 generations, allowing a detailed examination of inheritance.

#### Collection of naturally infesting families (F0 and F1 generations)

To obtain mites from the first two generations (F0 and F1), naturally infested families were collected from within the cells of 18- to 20-day-old worker bee pupae. At this developmental stage of the bee host, there is typically one dark brown female mite (the foundress, F0), one adult male (the foundress’ son, F1), at least one light-colored female (the first foundress’ daughter, F1), a few nymphs females, and an egg^25^. We determined each mite’s sex and developmental stage according to external morphology and cuticle coloring based on^50^. For the color index, see Fig S1. To ensure that all offspring belonged to a single family, we thoroughly examined the cell content to confirm that it was infested by a single foundress mite and that no other adults were present. Except for the adult female daughters (F1), the rest of the family members were preserved in a separate microcentrifuge tube kept at −20 °C for later DNA extraction. The adult F1 daughters were easily distinguished from the F0 foundress mite by their lighter cuticle color (color index 0-2, see Fig S1), and were assumed to have already copulated with their brother, the F1 male, by the time of the collection^20^. Next, the F1 young adult females were prepared to be artificially introduced into a freshly capped 5^th^ instar larva cell to start their first reproductive cycle.

#### Incubation and dispersal phases of F1 female mites

Based on our and others’ observations^51^, the survival and fecundity of young adult mites increase if they stay for a few days on an adult nurse bee before starting their first reproductive cycle. This period is presumably imitating the “dispersal phase” (former called the “phoretic phase”), in which the adult female mite infests and feeds on an adult honey bee^51,52^. Therefore, the collected young adult females (F1 generation) spent three days on a nurse bee before infestation. However, during the experiment, we noticed that the very young adult daughter’s cuticle was light and fragile, and most of the mites did not survive when they were transferred directly from the pupae onto a nurse bee. Therefore, we let the young mites first feed on a white-eyed pupa in a gelatin capsule for one to four days (“incubation phase”, Fig 5B1) until their cuticle was sclerotized and darker (color index 3-4, see Fig S1). After this intermediate period, the mites were marked on their dorsal idiosoma (PC-1MR POSCA, Japan), and transferred onto an adult nurse bee in a cage for another three days, for the above-mentioned “dispersal phase” (Fig 5B2). In all incubation periods, the mites were kept in a controlled dark environment, imitating hive conditions (34-34.5 °C, 60-75% RH).

#### Artificial infestation of F1 female mites

After the artificial “dispersal phase”, the mites were brushed off the bees using a fine paint-brush following anesthesia by a 20-s exposure to CO_2_, 0.2mPa at 10 L/hour, and placed on a moist filter paper to avoid desiccation until the infestation procedure. The mites were then introduced to a worker bee cell, one mite per cell, as described by^53^ (Fig 5C). In detail, six hours before infestation, cells containing a 5^th^ instar larva (about to be sealed) were marked on an acetate sheet attached to a frame from a mite-free hive. The frame was then placed back into the hive so that the bees could seal the cells. Five hours later the frame was taken into a warm room (25-28 °C) for the F1 females to be introduced into the freshly sealed cells. For the artificial infestation, a precise incision (3-5 mm) was made at the margin of the cell-capping using a surgical blade (N 11). After the incision was made, the cell capping was slightly lifted, just enough for the mite to crawl under the capping and into the cell. The mite was then placed at the entrance of the cell using a moist fine paintbrush, with its head facing the cell opening. After the mite entered the cell, it moved under the larva to the cell bottom. The cap was then resealed by gently pushing it down using the back end of the paintbrush. The infested frame was then incubated in a dark controlled environment (34-34.5 °C, 60-75% RH) for ten days, during which the mites were reproducing (Fig 5D). During this period emerging bees were removed daily.

#### Collection of an artificially infested family (F2 generation)

Ten days after the F1 mites’ infestation, the cells were opened and their contents were examined (Fig 5E). The family members’ sex, developmental stage (adult, nymph, or egg), and viability, were recorded. To assure the parenthood of the offspring (F2), double-infested cells and unmarked mites were not included in the final analysis. In total, out of 78 successful infestations, 14 were double-infested by a non-marked mite. Each individual was snap-frozen in a separate microcentrifuge tube at −20 °C for later DNA extraction. In total, we collected 223 mites belonging to 30 families. Each family was composed of a foundress adult female (F0), her male and female adult offspring (F1), and their F2 offspring (at least one male/female mite in various developmental stages).

### Genomic analysis of the Varroa pedigree

#### DNA extraction, library preparation, and sequencing

Genomic DNA was extracted from each mite following the protocol described previously^23^. In brief, after surface sterilization using absolute ethanol, each mite was crushed in a 1.5 mL microcentrifuge tube dipped in liquid nitrogen. The crushed mite was then processed using a QIAamp DNA Micro Kit (© Qiagen) following the manufacturer’s instructions and eluted in 20 μL of water. Total dsDNA quantity was measured using a Qubit™ 4 Fluorometer with an Invitrogen dsDNA HS Assay Kit, and DNA quality was verified using a Bioanalyzer 2100 with high-sensitivity DNA kits (Agilent, Japan). Short inserts of 150-bp paired-end libraries were prepared for each individual using a Nextera XT DNA Library Preparation Kit (Illumina, Japan) and 16 PCR cycles. cDNA was then cleaned and size-selected using CA-magnetic beads (Dynabeads® MyOne Carboxylic Acid, Invitrogen; Thermo Fisher Scientific, USA), and 11-11.5% PEG 6000 (Sigma-Aldrich, Japan). The library’s quality was measured using a Qubit™ 4 Fluorometer with an Invitrogen dsDNA HS Assay Kit, and its size was assessed using a 4200 Tapestation with a high-sensitivity D5000 kit (Agilent, Japan). Libraries were run on a NovaSeq6000 in 150 bp x 2 paired-end mode (Illumina, Japan) at the OIST Sequencing Center. All biosamples are available in the Sequence Read Archive (SRA) under the accession number PRJNA794941. Biosample details including sex, developmental stage, pedigree information, and sequencing coverage, are provided in Table S2.

#### Read mapping, variant calling, and filtering, of genomic data

We mapped all reads to the *V. destructor* reference genome GCF_002443255.1^54^ using the very sensitive mode of NextGenMap v0.5.0^55^. Reads were sorted and duplicates were removed using SAMtools^56^. We called variants against the reference genome using FreeBayes v1.1.0^57^ with the following parameters: minimum alternative allele counting = 5, minimum alternative allele fraction = 0.2, minimum mapping quality = 8, minimum base quality = 5, and using the four best SNP alleles. The variant calling resulted in a total of 73,903,468 variated sites. As our population consisted of closely related individuals (mite families from the same apiary, in a high-inbreeding system), we did not expect to detect many variants. Therefore, we set the minor allele frequency threshold to 0.2, which includes sites with a minor allele frequency of 20% or more, to exclude any rare variants due to mutations or possible mapping errors. We included variants only for the seven chromosomes (excluding the mitochondrial DNA and unscaffolded regions). In addition, we filtered for the following site quality parameters: we included only bi-allelic sites, a depth range between 16-40, and maximum missing data per site = 0.5. The variants’ quality distribution showed a clear bimodal distribution with a cutoff at ∼10,000 (Fig S2). To ensure the quality of the variants for the genetic inheritance analysis, we chose a strict minimum quality filter of 15,000. Filters were applied using VCFtools v0.1.12b^58^, resulting in 33,925 final variants from 223 individuals. The pipeline from fastq files down to the VCF file was constructed using Snakemake^59^. Codes are available and reproducible on GitHub (https://github.com/nurit-eliash/Varroa-pedigree-study). For further security of site quality, we applied two additional site-quality parameters to account for biases in alternate allele representation and to ensure high-confidence genotype calls based on allele frequency and quality. Variant allele frequency (VAF) was calculated as *VAF = AO/DP*, where AO (Alternate Observations) represents the number of reads supporting the alternate allele, and DP (Depth) is the total read depth at the site. Similarly, Quality Allele Frequency (QAF) was computed as *QAF = QA/QA+QO*, where QA (Quality of Alternate Allele) is the sum of Phred-scaled base quality scores for reads supporting the alternate allele, and QO (Quality of Reference Allele) is the sum of Phred-scaled base quality scores for reads supporting the reference allele. To minimize errors in genotyping due to sequencing artifacts or depth-related biases, we applied genotype-specific thresholds for VAF and QAF based on observation of values from F2 females of which parents and grandmother genotypes are fixed, and so their own genotype is assumed with high confidence (Fig S3, S4 and S5 for QAF and VAF distribution, in genotypes AA, BB and AB, in accordance). The thresholds (Table 1), ensure that homozygous reference (AA) sites exhibit low alternate allele support (VAF and QAF < 0.01), heterozygous (AB) sites maintain a balanced allele distribution (0.2< VAF < 0.8, 0.2 < QAF < 0.8), and homozygous alternate (BB) sites demonstrate high alternate allele support (VAF and QAF > 0.9). This filtering approach enhances the accuracy of genotype classification, while reducing the inclusion of low-confidence variant calls.

**Table 1.**
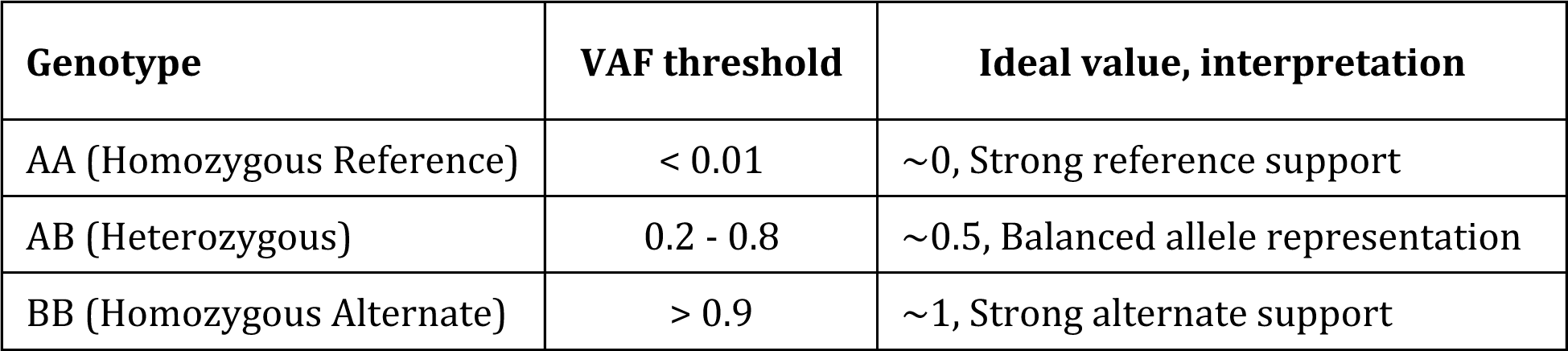
Quality Allele Frequency (QAF) and Variant allele frequency (VAF) threshold and their ideal values used for sites’ filtration in the genotype flow of mite pedigrees.

### Genotype validation using Sanger sequencing

Several genotype calls obtained from whole genome sequencing were validated by Sanger sequencing using the same genomic DNA extracted. For the validation, we randomly selected three genes, located on different chromosomes, each containing at least one variant site. In total, we re-sequenced 14 individual mites from four families on six variant sites. The sequenced region was located on the exons of three genes: alpha-mannosidase 2x-like (LOC111243621 on chromosome NW_019211455.1), 5-oxoprolinase-like (LOC111252938, on chromosome NW_019211459.1), and probable E3 ubiquitin-protein ligase RNF144A-A (LOC111254119, on chromosome NW_019211460.1). Primers were designed with the NCBI primer design tool (utilizing Primer3 and BLAST), with default parameters and product size set to 300-1,000 bp. Primer sequence, product length, and gene accession numbers are provided in Table S3. The size of the PCR product was checked by running it in 1% agarose gel (135V, 20 min), purified, and size selected using the solid phase reverse immobilization (SPRI) method. The purified PCR products were Sanger sequenced on a ThermoFisher SeqStudio Genetic Analyzer using the original reverse primer and BigDye® Direct Cycle Sequencing kit (Thermo Fisher Scientific, USA), following the manufacturer’s instructions. We then aligned the reverse complementary sequences to the Varroa reference genome (GCF_002443255.2_Vdes_3.0) to detect and validate the variant sites using Geneious Prime 2022.1.1. (https://www.geneious.com). For sequence alignments and the validated sites, see Fig S6.

### Transcriptomic analysis of Varroa mother and son

#### RNA extraction, library preparation, and sequencing

Total RNA was extracted from each single mite and sequenced by Metware Biotechnology Co., Ltd. (Wuhan, China) following an in-house protocol. RNA integrity was assessed using a Qubit™ 4 Fluorometer (Invitrogen; Thermo Fisher Scientific, USA) and a Qsep400 Bioanalyzer. For mRNA enrichment, polyadenylated RNA was isolated using Oligo(dT) beads, followed by fragmentation with fragmentation buffer. First-strand cDNA was synthesized using random hexamers and reverse transcriptase, while second-strand synthesis was carried out with dNTPs and DNA polymerase I. The resulting double-stranded cDNA was purified, subjected to end-repair, A-tailing, and adapter ligation, followed by PCR amplification. Library quality was assessed using a Qubit™ 4 Fluorometer with an Invitrogen dsDNA HS Assay Kit, and insert size distribution was confirmed using a Bioanalyzer (Agilent, Japan). The final libraries were pooled and sequenced on an Illumina platform in paired-end mode. All biosamples are available in the Sequence Read Archive (SRA) under the accession number PRJNA1216898. Biosample details including sex, family, and mapped reads, are provided in Table S1.

#### Read mapping, variant calling, and filtering, of RNAseq data

RNA-seq reads were mapped to the *V. destructor* reference genome (GCF_002443255.2_Vdes_3.0) using HISAT2 v2.2.1 in default settings^60^. The mapping rate varied between samples, with uniquely mapped reads ranging from ∼5.88% to 15.71% of the total reads. Clean reads were obtained after quality filtering using fastq v0.23.2^60^, with adapters removed, reads containing more than 10% N bases discarded, and bases with quality scores below 20 trimmed. Variant calling was performed using GATK ^62^, and SNPs and InDels were annotated with ANNOVAR^41^. To ensure high-confidence variant sites, we filtered out low-quality SNPs and retained only sites with adequate coverage and quality metrics, although the exact thresholds were not specified in the company report. We observed thousands of exonic variants per sample, including both synonymous and nonsynonymous SNVs, as well as stop-gain mutations. The resulting variant dataset included annotations for exonic, 5’ and 3’ UTR regions.

### Mathematical modeling

We followed the classical approach used by Wright^63^ to mathematically model declines in heterozygosity due to inbreeding within haplodiploid systems. This approach involves constructing path diagrams representing genetic transmission paths from ancestors to descendants and quantifying genetic relatedness through path coefficients. By enumerating paths and calculating their coefficients, we derived two mathematical relationships describing the expected declines in heterozygosity per generation under the reproductive life cycle of Varroa, one for the reproductive system with arrhenotokous diploid males and the other for haploid males. These two equations allowed a direct comparison between the two reproductive systems. Figure 3 plots the two equations in terms of how the rate of decline in heterozygosity, translated into effective population size, changes with the degree of inbreeding, which is measured by the probability of co-founding by females of the same mother. Mathematical details are in Supporting Information 1.

### Statistical analysis

All analyses were carried out in the R statistical environment^31^. Bioinformatic analyses and figures are available and reproducible directly from the online supplementary data (https://rpubs.com/Nurit_Eliash/1336154).

## Supporting information

Supplementary figures, tables and modeling math

## Acknowledgments

ASM was supported by a Future Fellowship from the Australian Research Council (FT160100178). HZ was supported by the earmarked fund for CARS (CARS-44). We wish to thank local beekeepers of Okinawa, Japan: Tsuto Arakaki, Takashi Ikemiya, and Ryo Kirino, from the Agricultural Environment Coordinator in Onna-village, for their help in providing Varroa-infested colonies, and Ms. Saori Chappell, OIST, for coordinating the sample collection and administrative helping throughout. At OIST, we thank Jan Moran and the HPC team for their support. We wish to thank Alison McAfee for providing the beautiful illustrations of Varroa mites and honey bees incorporated in the manuscript figures. We also thank Ms. Cristina de Morais Cianci from the Editorial Office of Genetics and Molecular Biology journal, for kindly providing a copy of the fundamental research on Varroa cytogenetics^31^. We also wish to thank Yoav Ram for commenting on the earlier draft of the manuscript. Finally, we wish to thank Mikheyev and Economo laboratory members for their continuous support and discussion throughout.

## References

1. Reed, D. H. & Frankham, R. Correlation between fitness and genetic diversity. Conserv. Biol. 17, 230–237 (2003).

2. Kliman, R., Sheehy, B. & Schultz, J. Genetic drift and effective population size. Nature Education 1, 3 (2008).

3. Papkou, A., Gokhale, C. S., Traulsen, A. & Schulenburg, H. Host-parasite coevolution: why changing population size matters. Zoology (Jena) 119, 330–338 (2016).

4. Frankham, R. & Ralls, K. Inbreeding leads to extinction: Conservation biology. Nature 392, 441–442 (1998).

5. Spielman, D., Brook, B. W., Briscoe, D. A. & Frankham, R. Does inbreeding and loss of genetic diversity decrease disease resistance? Conserv. Genet. 5, 439–448 (2004).

6. Charlesworth, D. & Willis, J. H. The genetics of inbreeding depression. Nat. Rev. Genet. 10, 783–796 (2009).

7. Venette, R. C. & Hutchison, W. D. Invasive insect species: Global challenges, strategies & opportunities. *Front*. Insect Sci. 1, 650520 (2021).

8. Siddiqui, J. A. et al. Insights into insecticide-resistance mechanisms in invasive species: Challenges and control strategies. Front. Physiol. 13, 1112278 (2022).

9. Allendorf, F. W. & Lundquist, L. L. Introduction: Population biology, evolution, and control of invasive species. Conserv. Biol. 17, 24–30 (2003).

10. Frankham, R. Resolving the genetic paradox in invasive species. Heredity 94, 385 (2005).

11. Sakai, A. K. et al. The population biology of invasive species. Annu. Rev. Ecol. Syst. 32, 305–332 (2001).

12. Suehs, C. M., Affre, L. & Médail, F. Invasion dynamics of two alien *Carpobrotus* (Aizoaceae) taxa on a Mediterranean island: II. Reproductive strategies. Heredity (Edinb.) 92, 550–556 (2004).

13. Manfredini, F., Arbetman, M. & Toth, A. L. A potential role for phenotypic plasticity in invasions and declines of social insects. Front. Ecol. Evol. 7, (2019).

14. Chevin, L. M. & Hoffmann, A. A. Evolution of phenotypic plasticity in extreme environments. Philos. Trans. R. Soc. Lond. B Biol. Sci. 372, (2017).

15. Tonione, M. A., Reeder, N. & Moritz, C. C. High genetic diversity despite the potential for stepping-stone colonizations in an invasive species of gecko on Moorea, French Polynesia. PLoS One 6, e26874 (2011).

16. Miyakawa, M. O. & Mikheyev, A. S. QTL mapping of sex determination loci supports an ancient pathway in ants and honey bees. PLoS Genet. 11, e1005656 (2015).

17. Rey, O., Facon, B., Foucaud, J., Loiseau, A. & Estoup, A. Androgenesis is a maternal trait in the invasive ant *Wasmannia auropunctata*. Proc. Biol. Sci. 280, 20131181 (2013).

18. Techer, M. A. et al. Divergent evolutionary trajectories following speciation in two ectoparasitic honey bee mites. Communications Biology 2, 357 (2019).

19. Frey, E., Schnell, H. & Rosenkranz, P. Invasion of *Varroa destructor* mites into mite-free honey bee colonies under the controlled conditions of a military training area. J. Apic. Res. 50, 138–144 (2011).

20. Häußermann, C. K. et al. Reproductive parameters of female Varroa destructor and the impact of mating in worker brood of Apis mellifera. Apidologie 51, 342–355 (2020).

21. Oldroyd, B. P. Coevolution while you wait: *Varroa jacobsoni*, a new parasite of western honeybees. Trends Ecol. Evol. 14, 312–315 (1999).

22. Dietemann, V. et al. Population genetics of ectoparasitic mites *Varroa* spp. in Eastern and Western honey bees. Parasitology 1–11 (2019).

23. Techer, M. A., Roberts, J. M. K., Cartwright, R. A. & Mikheyev, A. S. The first steps toward a global pandemic: Reconstructing the demographic history of parasite host switches in its native range. Molecular ecology 31, 1358–1374 (2022).

24. Mitton, G. A. et al. More than sixty years living with *Varroa destructor*: a review of acaricide resistance. Int. J. Pest Manage. 1–18 (2022).

25. Rosenkranz, P., Aumeier, P. & Ziegelmann, B. Biology and control of *Varroa destructor*. J. Invertebr. Pathol. 103, 96–119 (2010).

26. Traynor, K. S., et al. *Varroa destructor*: A complex parasite, crippling honey bees worldwide. Trends Parasitol. 36, 592–606 (2020).

27. Nazzi, F. & Le Conte, Y. Ecology of *Varroa destructor*, the major ectoparasite of the western honey bee, *Apis mellifera*. Annu. Rev. Entomol. 61, 417–432 (2016).

28. Metcalf, R. A., Marlin, J. C. & Whitt, G. S. Low levels of genetic heterozygosity in hymenoptera. Nature 257, 792–794 (1975).

29. Moro, A. et al. Adaptive population structure shifts in invasive parasitic mites, *Varroa destructor*. Ecol. Evol. 11, 5937–5949 (2021).

30. Tien, N. S. H., Sabelis, M. W. & Egas, M. Inbreeding depression and purging in a haplodiploid: gender-related effects. Heredity (Edinb.) 114, 327–332 (2015).

31. Steiner, J., Pompolo, S. D., Takahashi, C. S. & Goncalves, L. S. Cytogenetics of the acarid *Varroa jacobsoni*. Rev. Bras. Genet. 4, 841–844 (1982).

32. Ruijter, A. & Pappas, N. Karyotype and sex determination of *Varroa jacobsoni* Oud. in Varroa jacobsoni Oud. affecting honey bees: present status and needs (ed. Cavalloro, R.) 41–44 (Rotterdam: Published for the Commission of the European Communities, 1983).

33. Bachtrog, D. et al. Sex determination: why so many ways of doing it? PLoS Biol. 12, e1001899 (2014).

34. Bull, J. J. Sex determining mechanisms: an evolutionary perspective. Experientia 41, 1285–1296 (1985).

35. Cruickshank, R. H. & Thomas, R. H. Evolution of haplodiploidy in *dermanyssine* mites (Acari: Mesostigmata). Evolution 53, 1796–1803 (1999).

36. Blackmon, H., Hardy, N. B. & Ross, L. The evolutionary dynamics of haplodiploidy: Genome architecture and haploid viability. Evolution 69, 2971–2978 (2015).

37. Oh, J. et al. Molecular phylogeny reveals Varroa mites are not a separate family but a subfamily of Laelapidae. Sci. Rep. 14, 13994 (2024).

38. Tree of Sex Consortium. Tree of Sex: a database of sexual systems. Sci. Data 1, 140015 (2014).

39. DE Moraes, G. J., et al. Catalogue of the free-living and arthropod-associated *Laelapidae canestrini* (Acari: Mesostigmata), with revised generic concepts and a key to genera. Zootaxa 5184, 1–509 (2022).

40. Normark, B. B. The evolution of alternative genetic systems in insects. Annu. Rev. Entomol. 48, 397–423 (2003).

41. Wright, S. Inbreeding and homozygosis. Proc. Natl. Acad. Sci. U. S. A. 19, 411–420 (1933).

42. Ross, L., Mongue, A. J., Hodson, C. N. & Schwander, T. Asymmetric inheritance: The diversity and Evolution of non-Mendelian reproductive strategies. Annu. Rev. Ecol. Evol. Syst. 53, 1–23 (2022).

43. Forni, G., Mantovani, B., Mikheyev, A. S. & Luchetti, A. Parthenogenetic stick insects exhibit signatures of preservation in the molecular architecture of male reproduction. Genome Biol. Evol. 16, (2024).

44. Bull, J. J. An advantage for the evolution of male haploidy and systems with similar genetic transmission. Heredity 43, 361–381 (1979).

45. Neiman, M., Meirmans, S. & Meirmans, P. G. What can asexual lineage age tell us about the maintenance of sex? Ann. N. Y. Acad. Sci. 1168, 185–200 (2009).

46. Sasaki, K. & Obara, Y. Egg activation and timing of sperm acceptance by an egg in honeybees (*Apis mellifera* L.). Insectes Soc. 49, 234–240 (2002).

47. Moro, A., Blacquière, T., Panziera, D., Dietemann, V. & Neumann, P. Host-parasite co-evolution in real-time: Changes in honey bee resistance mechanisms and mite reproductive strategies. Insects 12, 120 (2021).

48. Faber, N. R., Meiborg, A. B., Mcfarlane, G. R., Gorjanc, G. & Harpur, B. A. A gene drive does not spread easily in populations of the honey bee parasite *Varroa destructor*. Apidologie 52, 1112–1127 (2021).

49. Beaurepaire, A. L. et al. Population genetics of ectoparasitic mites suggest arms race with honeybee hosts. Sci. Rep. 9, 11355 (2019).

50. Büchler, R., Costa, C., Mondet, F., Kezic, N. & Kovacic, M. Screening for Low Varroa Mite Reproduction (SMR) and Recapping in European Honey Bees. https://www.beebreeding.net/wp-content/uploads/2017/11/RNSBB_SMR-recapping_protocol_2017_09_11.pdf (2017).

51. Xie, X., Huang, Z. Y. & Zeng, Z. Why do Varroa mites prefer nurse bees? Sci. Rep. 6, 28228 (2016).

52. Piou, V., Tabart, J., Urrutia, V., Hemptinne, J.-L. & Vétillard, A. Impact of the phoretic phase on reproduction and damage caused by *Varroa destructor* (Anderson and Trueman) to its host, the European honey bee (*Apis mellifera* L.). PLoS One 11, e0153482 (2016).

53. Dietemann, V. et al. Standard methods for varroa research. J. Apic. Res. 52, 1–54 (2013).

54. Sedlazeck, F. J., Rescheneder, P. & von Haeseler, A. NextGenMap: fast and accurate read mapping in highly polymorphic genomes. Bioinformatics 29, 2790–2791 (2013).

55. Li, H. et al. The Sequence Alignment/Map format and SAMtools. Bioinformatics 25, 2078–2079 (2009).

56. Garrison, E. & Marth, G. Haplotype-based variant detection from short-read sequencing. arXiv [q-bio.GN] (2012).

57. Danecek, P. et al. Twelve years of SAMtools and BCFtools. Gigascience 10, giab008 (2021).

58. Köster, J. & Rahmann, S. Snakemake--a scalable bioinformatics workflow engine. Bioinformatics 28, 2520–2522 (2012).

59. Kim, D., Langmead, B. & Salzberg, S. L. HISAT: a fast spliced aligner with low memory requirements. Nat. Methods 12, 357–360 (2015).

60. Chen, S., Zhou, Y., Chen, Y. & Gu, J. fastp: an ultra-fast all-in-one FASTQ preprocessor. Bioinformatics 34, i884–i890 (2018).

61. McKenna, A. et al. The Genome Analysis Toolkit: a MapReduce framework for analyzing next-generation DNA sequencing data. Genome Res. 20, 1297–1303 (2010).

62. Wang, K., Li, M. & Hakonarson, H. ANNOVAR: functional annotation of genetic variants from high-throughput sequencing data. Nucleic Acids Res. 38, e164 (2010).

63. Team, R. C. R: A language and environment for statistical computing. (2013).

