## Supplementary figures, tables and modeling math for "Varroa mites escape the evolutionary trap of haplodiploidy"

**This PDF file includes:**

Supporting text, modeling mathematics S1
Figures S1 to S6
Tables S1 to S3
SI References

**Supporting Information Text**

**Modeling mathematics S1.** Path coefficient modelling for rate of decrease in heterozygosity.

We adapted the classical approach used by Wright [1] to mathematically model declines in heterozygosity $p$ (proportion of heterozygosis) due to inbreeding under the reproductive life cycle of *Varroa destructor*. In particular, we allow Varroa population size to change over generation and calculate the corresponding heterozygosity recursively under two reproductive systems. In both systems, females are diploid and produced sexually. In one system, mothers produce clone sons. In the other system, mothers produce haploid sons. We make the following assumptions to make the model analytical.

Assumptions:

1) The parasite population starts from one female, so $N(0) = 1$
2) The number of females changes deterministically following a logistics growth model, where population size at $t + 1$ generation is $N(t + 1) = \left[1 + r(1 - \frac{N(t)}{K})\right] N(t)$. In the equation, $K$ is carrying capacity, approximating the total number of blood cells where females can lay eggs; $r$ is intrinsic growth rate, approximated by $n - 1$, where $n$ is the number of daughters each female has.
3) Each female has $P$ probability of laying eggs in an empty blood cell and produces $n$ daughters and 1 son. Assuming females laying eggs randomly in $K$ blood cells, $P(t) = (1 - \frac{1}{K})^{N(t)-1}$
4) Male(s) and females in the same cell have equal chance of mating to produce daughters.

Using the notation in [1], Wright's approach starts with deriving correlation between uniting gametes in the current generation $f$ as a function of $f'$, $f''$, and so on, where the number of prime indicates how many generations ago.

To derive the equation for $f$, let's consider a daughter, whose father and mother may have the same mother ($F = M$) or different mothers ($F \neq M$). Under each case, we can calculate the probability of the case given the reproductive life cycle of Varroa and use

Path coefficient modelling to derive $f$ as a function of $f'$, $f''$, as well as $f_f{}'$ and $f_m{}'$, which are the correlation between gametes from different females and from different males in the previous generation. For example, under the haplodiploid system, the two cases are shown in the table below, where in the pedigree, circles are females, squares are males, and dots are gametes. Arrows shows how a gamete from an individual is passed on to the next generation with $b$ giving the correlation between the individual's genome and its gametes and $a$ giving the correlation between the gamete and the genome of the offspring that inherits the gamete. Path in different colours show different ways of how two gametes of the daughter in the current generation are related. The daughter in the current generation is sit at the bottom of the pedigree.

| Case | Pedigree | Correlation | Probability |
|---|---|---|---|
| $F = M$ | 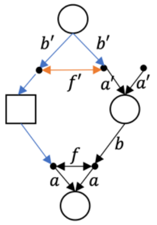  | $ba'(f' + b'^2) = \frac{1}{2}\left(\frac{1}{2} + f' + \frac{1}{2}f''\right)$ | $P + (1 - P)\frac{1}{2}$ |
| $F \neq M$ | 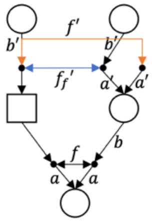  | $ba'(f' + f_f') = \frac{1}{2}(f' + f_f{}')$ | $(1 - P)\frac{1}{2}$ |

Summing these two cases together, we have

$$f = \frac{P+1}{2}\left(\frac{1}{4} + \frac{1}{4}f''\right) + \frac{1}{2}f' + \frac{1-P}{4}f_f{}' \qquad (1)$$

The initial condition is defined by $f(1)$, the correlation between gametes in the founder female. Generation 2 can only have father and mother from the same mother ($F = M$), which is confirmed by the probability term when $P = 1$ because $N(0) = 1$. Therefore, $f(2) = \frac{1}{2}\left(\frac{1}{2} + f(1) + \frac{1}{2}f(0)\right)$. Assuming that the founder female is from a population where correlation between gametes is stabilized, i.e., $f(1) = f(0)$, $f(2) = \frac{1}{2}\left(\frac{1}{2} + \frac{3}{2}f(1)\right)$.

In equation (1), $f$ is related to $f_f{}'$. To calculate $f_f{}'$, let's consider two daughters who share both father and mother ($F1 = F2; M1 = M2$), only share father ($F1 = F2; M1 \neq M2$), only share mother ($F1 \neq F2$; $M1 = M2$), and do not share either father or mother ( $F1 \neq F2$; $M1 \neq M2$). Again, under each case, we can calculate the probability of the case given the reproductive life cycle of Varroa and use Path coefficient modelling to derive $f_f$. For example, under the haplodiploid system, the four cases are shown in the table below.

| Case | Pedigree | Correlation | Probability |
|---|---|---|---|
| $F1 = F2$<br>$M1 = M2$ | 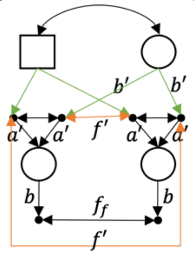 | $b^2a'^2(2f' + 1 + b'^2)$<br>$= \frac{1}{8}(3 + 4f' + f'')$ | $\frac{1}{4nN}(3P + 1)$ |
| $F1 = F2$<br>$M1 \neq M2$ | 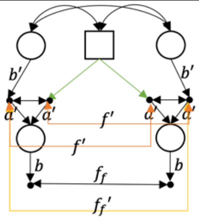 | $b^2a'^2\left(2f' + f_f' + 1\right)$<br>$= \frac{1}{4}\left(1 + 2f' + f_f'\right)$ | $\frac{(1+P)^2}{4N} - \frac{P}{nN}$ |
| $F1 \neq F2$<br>$M1 = M2$ | 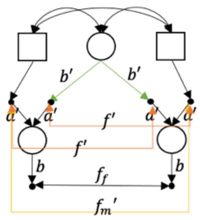 | $b^2a'^2(2f' + f_m' + b'^2) =$<br>$\frac{1}{4}\left(2f' + \left(1 - \frac{1}{N}\right)f_f'' + \frac{1}{2} + \frac{1}{2}f''\right)$ | $(1 - P)\frac{1}{nN}\frac{1}{4}$ |
| $F1 \neq F2$<br>$M1 \neq M2$ | 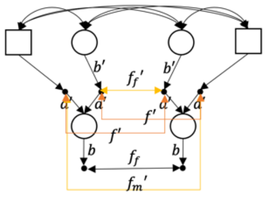 | $b^2a'^2\left(2f' + f_f' + f_m'\right)$<br>$= \frac{1}{4}\left(2f' + f_f' + \left(1 - \frac{1}{N}\right)f_f''\right)$ | $1 + \frac{P-1}{2nN}$<br>$- \frac{(1+P)^2}{4N}$ |

In this table, $f_m{}'$ is a function of $f_f'$. Since each mother only produces one son, two sons are always from different mothers ($F1 \neq F2$). Under the haplodiploid system, the case is shown in the table below, which gives us $f_m = \left(1 - \frac{1}{N}\right)f_f'$.

| Case | Pedigree | Correlation | Probability |
|---|---|---|---|
| $F1 \neq F2$ | 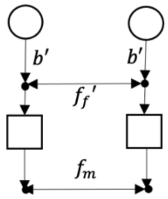  | $f_f'$ | $1-\frac{1}{N}$ |

Summing over the four cases for $f_f$, we have

$$\begin{aligned} f_f = &\frac{3P+1}{4nN}\frac{1}{8}(3+4f'+f'') + \left[\frac{(1+P)^2}{4N} - \frac{P}{nN}\right]\frac{1}{4}\left(1+2f'+f_f'\right) \\ &+ \frac{1-P}{4nN}\frac{1}{4}\left(2f' + \left(1-\frac{1}{N}\right)f_f'' + \frac{1}{2} + \frac{1}{2}f''\right) \\ &+ \left[1 + \frac{P-1}{2nN} - \frac{(1+P)^2}{4N}\right]\frac{1}{4}\left(2f' + f_f' + \left(1-\frac{1}{N}\right)f_f''\right) \end{aligned} \tag{2}$$

The initial condition is $f_f(1) = f(1) = f(0)$, because females in generation 1 must have the same mother who is the founder. Females in Generation 2 can only have the same father ($F1 = F2$). This is confirmed by the probability term when $N = 1$ and $P = 1$. As a result, $f_f(2) = \frac{1}{8n}[1 - f(1)] + \frac{1}{4}[1 + 3f(1)]$.

We can replace $f_f$, $f_f{}'$, $f_f{}''$ in equation (1) with functions of $f$, $f'$, $f''$, using equation (2). This gives us

$$\begin{aligned} f = &\frac{P+1}{2}\left(\frac{1}{4} + \frac{1}{4}f''\right) + \frac{1}{2}f' \\ &+ \frac{1-P}{4}\left\{\frac{3P+1}{4nN}\frac{1}{8}(3+4f''+f''') + \left[\frac{(1+P)^2}{4N} - \frac{P}{nN}\right]\frac{1}{4}(1+2f'')\right. \\ &+ \frac{1-P}{4nN}\frac{1}{4}\left(2f'' + \frac{1}{2} + \frac{1}{2}f'''\right) + \left[1 + \frac{P-1}{2nN} - \frac{(1+P)^2}{4N}\right]\frac{1}{2}f'' \\ &+ \left[-\frac{P}{nN} + 1 + \frac{P-1}{2nN}\right]\frac{1}{P-1}\left(\frac{P+1}{2}\left(\frac{1}{4} + \frac{1}{4}f'''\right) + \frac{1}{2}f'' - f'\right) \\ &+ \left[1 - \frac{1-P}{4nN} - \frac{(1+P)^2}{4N}\right]\left(1-\frac{1}{N}\right)\frac{1}{P-1}\left(\frac{P+1}{2}\left(\frac{1}{4} + \frac{1}{4}f''''\right) + \frac{1}{2}f''' \right. \\ &\left.\left. - f''\right)\right\} \end{aligned} \tag{3}$$

Rearrange equation (3) to the form of $f = c_0 + c_1 f' + c_2 f'' + c_3 f''' + c_4 f''''$, with initial conditions: $f(1) = f(0)$, $f(2) = \frac{1}{2}\left(\frac{1}{2} + \frac{3}{2} f(1)\right)$. For $f(3)$, we know $f_f(2) = \frac{1}{8n}[1 - f(1)] + \frac{1}{4}[1 + 3f(1)]$, so using equation (1), $f(3) = \frac{P+1}{2}\left(\frac{1}{4} + \frac{1}{4} f(1)\right) + \frac{1}{2} f(2) + \frac{1-P}{4} f_f(2)$, where $P = (1 - \frac{1}{K})^{N(1)-1}$.

Substituting $f = 1 - \frac{p}{p_0}$ in equation (3), where $p_0$ is the proportion of heterozygosis under random mating, we have

$$1 - \frac{p}{p_0} = c_0 + c_1\left(1 - \frac{p'}{p_0}\right) + c_2\left(1 - \frac{p''}{p_0}\right) + c_3\left(1 - \frac{p'''}{p_0}\right) + c_4\left(1 - \frac{p''''}{p_0}\right)$$

Since $c_0 + c_1 + c_2 + c_3 + c_4 = 1$

$$p = c_1 p' + c_2 p'' + c_3 p''' + c_4 p''''$$

with initial conditions: $p(t) = [1 - f(t)]p_0$, where $t = 0,1,2,3$.

The loss rate in heterozygosity at generation $t$ is defined as $\frac{p(t)}{p(t-1)} - 1$.

Now let's repeat the derivation for the system where mothers produce clone sons. The two cases for $f$ are summarised below.

| Case | Pedigree | Correlation | Probability |
|---|---|---|---|
| $F = M$ | 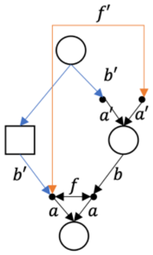 | $ba'(f' + b'^2) = \frac{1}{2}\left(\frac{1}{2} + f' + \frac{1}{2} f''\right)$ | $P + (1 - P)\frac{1}{2}$ |

| | | | |
|---|---|---|---|
| $F \neq M$ | 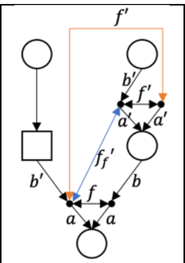 | $ba'(f' + f_f') = \frac{1}{2}(f' + f_f{}')$ | $(1-P)\frac{1}{2}$ |

The main difference to having haploid sons is that now meiosis ($b'$) happens when the son produces daughter, rather than when the mother produces the son. This difference does not affect the relationship between $f$ and $f_f{}'$ as the functions in the table above is the same as those in the corresponding table for having haploid sons.

However, the difference affects $f_f$, for which the four cases are summarised below. Intuitively, $f_m$ is unrelated here, because all males are the same as their mothers.

| Case | Diagram | Correlation | Probability |
|---|---|---|---|
| $F1 = F2$ $M1 = M2$ | 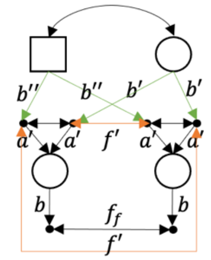 | $b^2a'^2(2f' + b''^2 + b'^2) = \frac{1}{4}\left(1 + 2f' + \frac{1}{2}f'' + \frac{1}{2}f'''\right)$ | $\frac{1}{4nN}(3P+1)$ |
| $F1 = F2$ $M1 \neq M2$ | 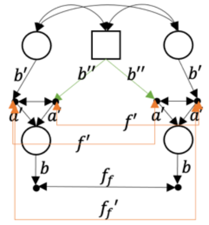 | $b^2a'^2(2f' + f_f' + b''^2) = \frac{1}{4}\left(\frac{1}{2} + 2f' + \frac{1}{2}f''' + f_f'\right)$ | $\frac{(1+P)^2}{4N} - \frac{P}{nN}$ |
| $F1 \neq F2$ $M1 = M2$ | 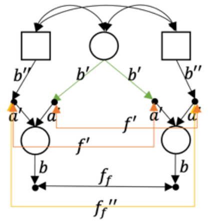 | $b^2a'^2(2f' + f_f'' + b'^2) = \frac{1}{4}\left(\frac{1}{2} + 2f' + \frac{1}{2}f'' + f_f''\right)$ | $(1-P)\frac{1}{nN}\frac{1}{4}$ |

| | | | |
|---|---|---|---|
| $F1 \neq F2$ <br> $M1 \neq M2$ | 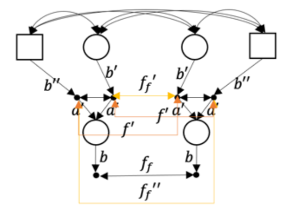 | $b^2 a'^2 \left(2f' + f_f' + f_f''\right)$ <br> $= \frac{1}{4}\left(2f' + f_f' + f_f''\right)$ | $1 + \frac{P-1}{2nN} - \frac{(1+P)^2}{4N}$ |

Summing the four cases together, we have

$$f_f = \frac{3P+1}{4nN}\frac{1}{4}\left(1 + 2f' + \frac{1}{2}f'' + \frac{1}{2}f'''\right) + \left[\frac{(1+P)^2}{4N} - \frac{P}{nN}\right]\frac{1}{4}\left(\frac{1}{2} + 2f' + \frac{1}{2}f''' + f_f'\right)$$
$$+ \frac{1-P}{4nN}\frac{1}{4}\left(\frac{1}{2} + 2f' + \frac{1}{2}f'' + f_f''\right) + \left[1 + \frac{P-1}{2nN} - \frac{(1+P)^2}{4N}\right]\frac{1}{4}\left(2f' + f_f' + f_f''\right)$$

The initial condition is $f_f(1) = f(1) = f(0) = f(-1)$. Females in Generation 2 can only have the same father ($F1 = F2$), so $f_f(2) = \frac{1}{8n}[1 - f(1)] + \frac{1}{8}[1 + 7f(1)]$.

Substituting $f_f, f_f', f_f''$ using equation 1,

$$f = \frac{P+1}{2}\left(\frac{1}{4} + \frac{1}{4}f''\right) + \frac{1}{2}f'$$
$$+ \frac{1-P}{4}\left\{\frac{3P+1}{4nN}\frac{1}{4}\left(1 + 2f'' + \frac{1}{2}f''' + \frac{1}{2}f''''\right)\right.$$
$$+ \left[\frac{(1+P)^2}{4N} - \frac{P}{nN}\right]\frac{1}{4}\left(\frac{1}{2} + 2f'' + \frac{1}{2}f''''\right) + \frac{1-P}{4nN}\frac{1}{4}\left(\frac{1}{2} + 2f'' + \frac{1}{2}f'''\right)$$
$$+ \left[1 + \frac{P-1}{2nN} - \frac{(1+P)^2}{4N}\right]\frac{1}{4}2f''$$
$$+ \left[1 - \frac{P+1}{2nN}\right]\frac{1}{P-1}\left(\frac{P+1}{2}\left(\frac{1}{4} + \frac{1}{4}f'''\right) + \frac{1}{2}f'' - f'\right)$$
$$\left.+ \left[1 + \frac{P-1}{4nN} - \frac{(1+P)^2}{4N}\right]\frac{1}{P-1}\left(\frac{P+1}{2}\left(\frac{1}{4} + \frac{1}{4}f''''\right) + \frac{1}{2}f''' - f''\right)\right\}$$

with initial conditions: $f(1) = f(0)$, $f(2) = \frac{1}{2}\left(\frac{1}{2} + \frac{3}{2}f(1)\right)$. For $f(3)$, we know $\frac{1}{8n}[1 - f(1)] + \frac{1}{8}[1 + 7f(1)]$, so using equation (1), $f(3) = \frac{P+1}{2}\left(\frac{1}{4} + \frac{1}{4}f(1)\right) + \frac{1}{2}f(2) + \frac{1-P}{4}f_f(2)$, where $P = (1 - \frac{1}{K})^{N(1)-1}$.

The rest of the derivation is the same.

**Supporting figures**

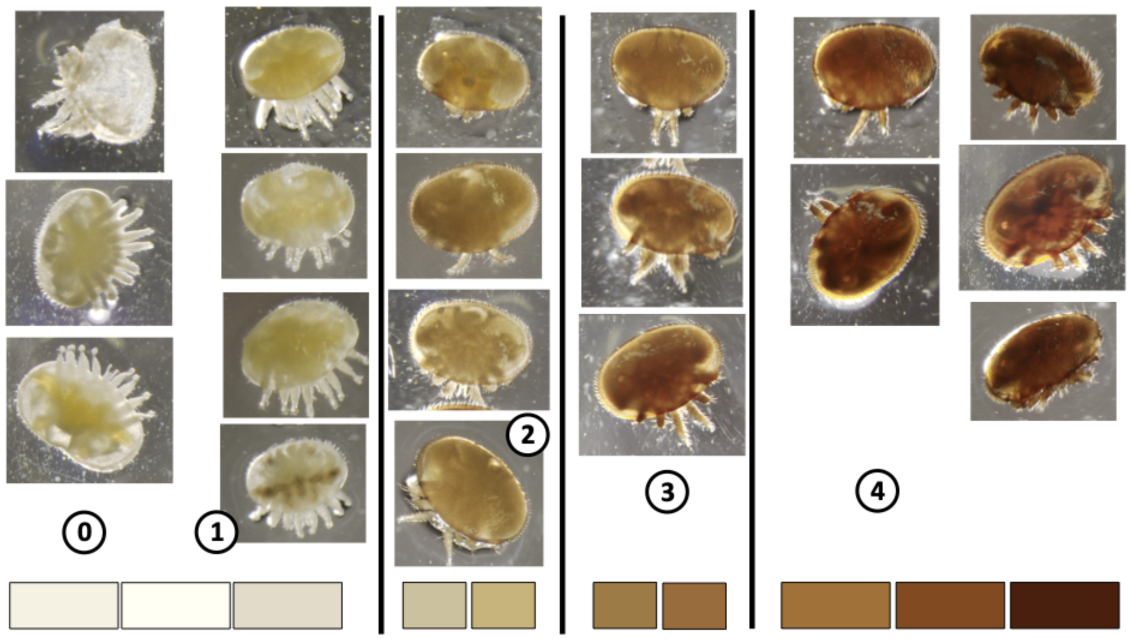

**Figure S1.** The color scales of Varroa adult female mites help differentiate the mother from her daughters. Mites were colored from light white (0) to dark brown (4). Adult female mites in the color of (4) were considered Foundress mites and were used as F0 generation in the artificial infestation experiment. Adult female mites in the colours between 2 and 3 were considered adult daughters of F0 and were used as the F1 generation. Photographs by Maeva Techer.

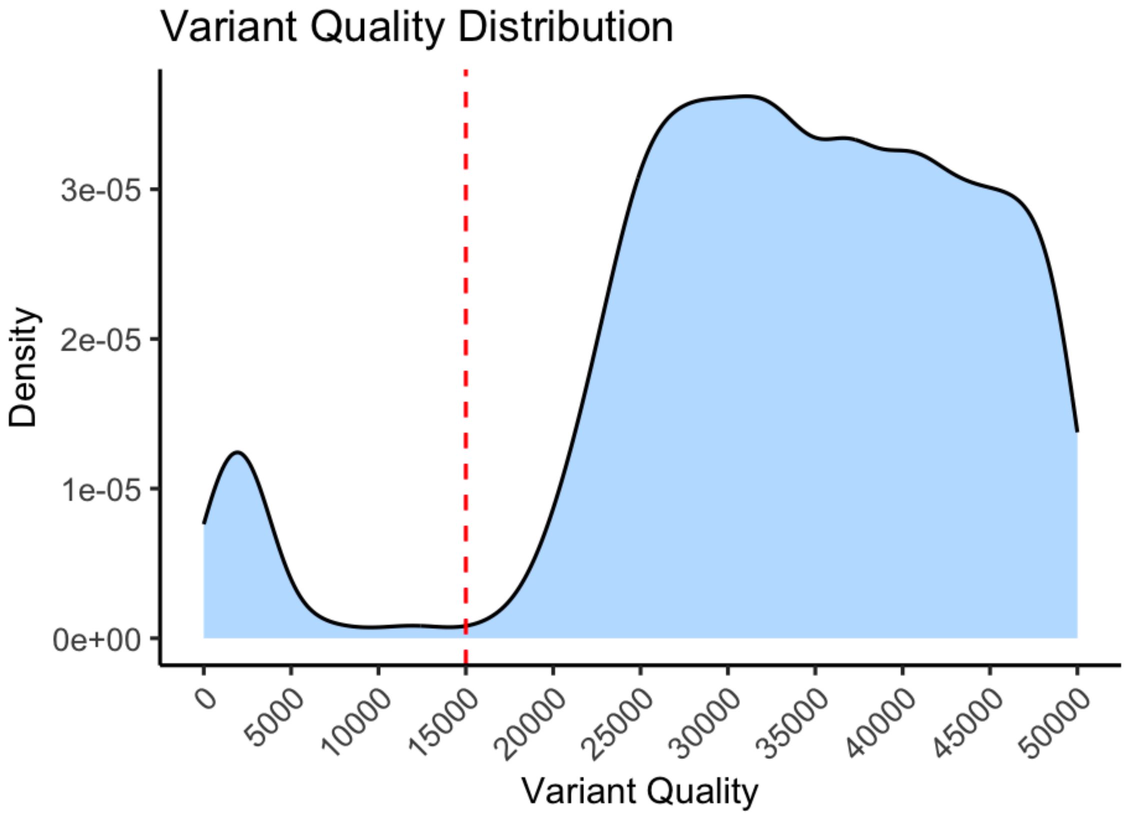

**Figure S2.** Quality distribution of variants filtered for a minimum quality of 40. The plot shows a clear bimodal distribution with a cutoff at ~10,000. To ensure the quality of the variants for the genetic inheritance analysis, we chose a strict minimum quality filter of 15,000.

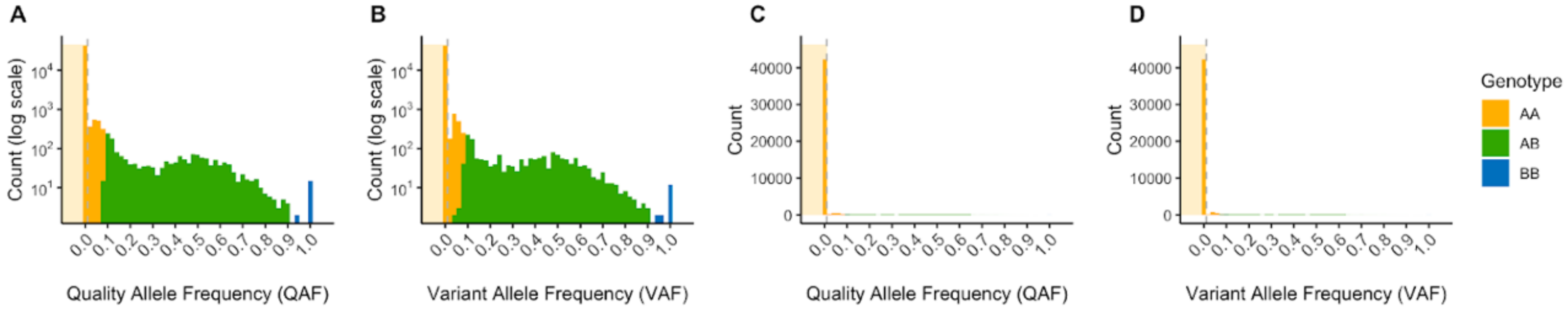

**Figure S3. Distribution of Quality Allele Frequency (QAF) and Variant Allele Frequency (VAF) values in F2 females from crosses with homozygous AA parents. A-B. QAF and VAF distributions shown on logarithmic scales. C-D. Same distributions shown on linear scales.** In all plots, shaded regions indicate the thresholds for genotype classification: AA (yellow, < 0.01). Vertical dashed lines mark the threshold boundaries. The plots show the distribution of genotype calls, with AA genotypes predominantly falling below the 0.01 threshold range.

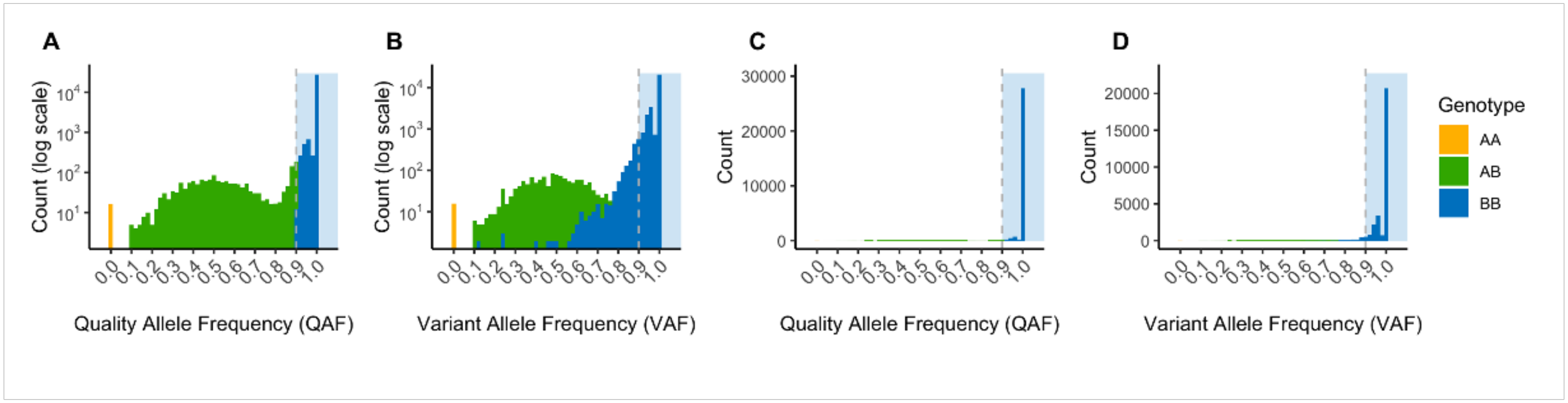

**Figure S4. Distribution of Quality Allele Frequency (QAF) and Variant Allele Frequency (VAF) values in F2 females from crosses with homozygous BB parents. A-B. QAF and VAF distributions shown on logarithmic scales. C-D. Same distributions shown on linear scales.** In all plots, shaded regions indicate the thresholds for genotype classification: BB (blue, > 0.9). Vertical dashed lines mark the threshold boundaries. The plots show the distribution of genotype calls, with BB genotypes predominantly falling above the 0.9 threshold range.

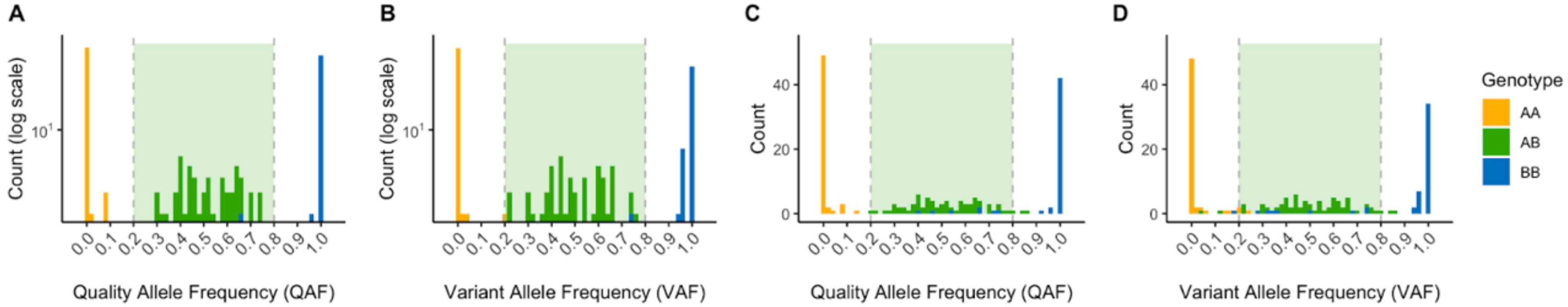

**Figure S5. Distribution of Quality Allele Frequency (QAF) and Variant Allele Frequency (VAF) values in F2 females from crosses with heterozygous AB parents. A-B. QAF and VAF distributions shown on logarithmic scales. C-D. Same distributions shown on linear scales.** In all plots, shaded regions indicate the thresholds for genotype classification: AB (green, 0.2-0.8). Vertical dashed lines mark the threshold boundaries. The plots show the distribution of genotype calls, with AB genotypes predominantly falling within the 0.2-0.8 threshold range.

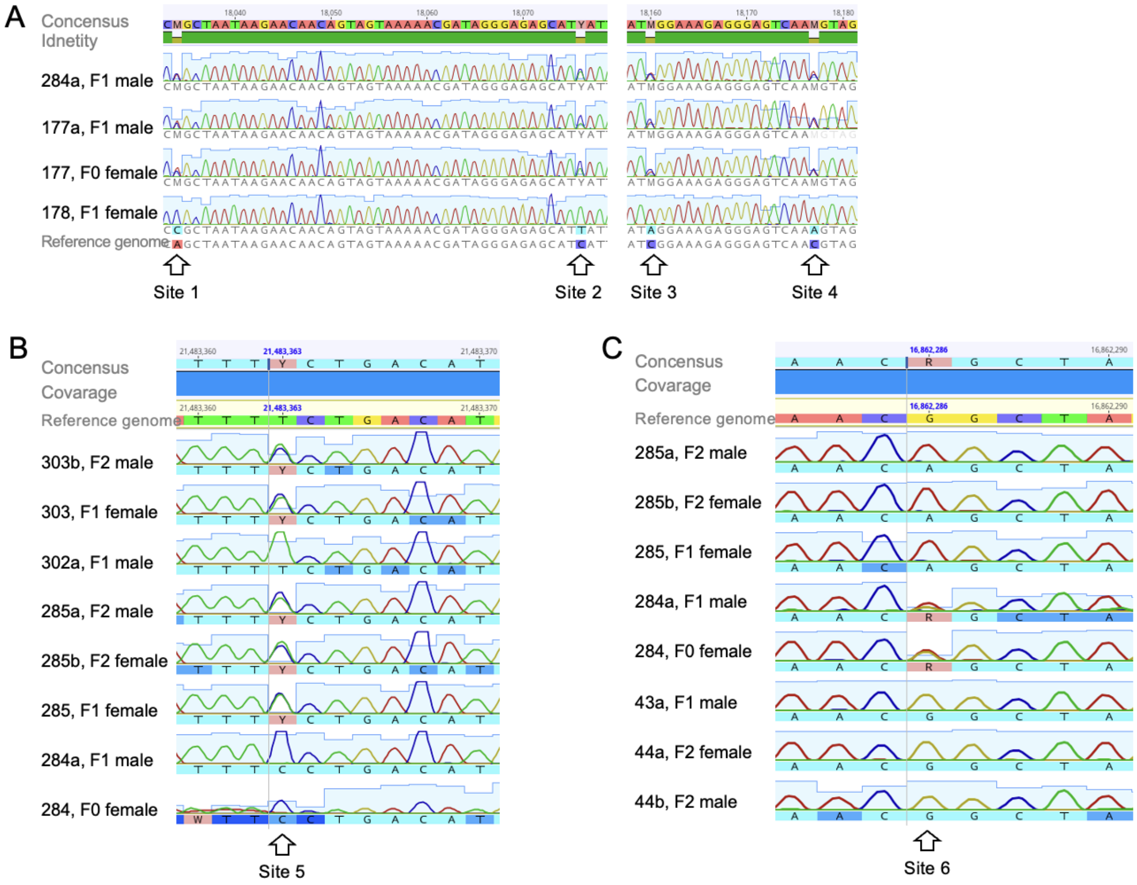

**Figure S6.** Alignment of Sanger-sequenced regions amplified using mite DNA from different families as a template. The amplicon sequences were aligned to a Varroa mite gene from the reference genome. (A) Amplicons of alpha-mannosidase 2x-like gene (LOC111243621), amplified on the DNA template of two adult males (284a, 177a) and two adult females (177 and her daughter 178); (B) Amplicons of 5-oxoprolinase-like gene (LOC111252938), amplified on the DNA template of family 302, and family 284; (C) Amplicons of probable E3 ubiquitin-protein ligase RNF144A-A gene (LOC111254119), amplified on the DNA template of family 284 and 43.

### Supporting tables

**Table S1. Detailed information for each of the Varroa mite samples sequenced for transcriptomic analysis**. All samples belong to the *Varroa destructor* mite collected from *Apis mellifera* colonies at Zhejiang University, China, in November 2024. RNA was extracted from whole mite bodies, and RNA libraries were constructed and sequenced on the Illumina platform by Metware Biotechnology Co., Ltd. (Wuhan, China). The sequencing includes an adult female mite and her adult male son from five different families. Reads were mapped to the Varroa destructor reference genome (GCF_002443255.2_Vdes_3.0) using HISAT2.

| Sample name | Sample species | Host species | Family | Developmental stage | Sex | Mapped reads |
|---|---|---|---|---|---|---|
| M-5 | *Varroa destructor* | *Apis mellifera* | 5 | Adult | Male | 8,906,226 |
| M-4 | *Varroa destructor* | *Apis mellifera* | 4 | Adult | Male | 7,953,816 |
| M-3 | *Varroa destructor* | *Apis mellifera* | 3 | Adult | Male | 6,620,283 |
| M-2 | *Varroa destructor* | *Apis mellifera* | 2 | Adult | Male | 9,012,902 |
| M-1 | *Varroa destructor* | *Apis mellifera* | 1 | Adult | Male | 9,238,715 |
| FF-5 | *Varroa destructor* | *Apis mellifera* | 5 | Adult | Female | 6,708,926 |
| FF-4 | *Varroa destructor* | *Apis mellifera* | 4 | Adult | Female | 6,567,500 |
| FF-3 | *Varroa destructor* | *Apis mellifera* | 3 | Adult | Female | 5,436,066 |
| FF-2 | *Varroa destructor* | *Apis mellifera* | 2 | Adult | Female | 3,728,485 |
| FF-1 | *Varroa destructor* | *Apis mellifera* | 1 | Adult | Female | 6,730,915 |

**Table S2.** Detailed information for each of the Varroa mites sequenced for the pedigree construction. All samples belong to *Varroa destructor* mite, collected from a Western honey bee (*Apis mellifera* L.) sealed cell from a hive located at OIST experimental apiary, Onna-son Okinawa, Japan. Samples were collected by Nurit Eliash from July to October 2020, and extracted from whole-body tissue by Endo Tatsuya at OIST.

| | | | | | | | |
|---|---|---|---|---|---|---|---|
| 302_303b_grnson | SAMN24905542 | 302 | Adult | 2 | male | 98.94% | 23.9 |
| 322_322_fnd | SAMN24905543 | 322 | Adult | 0 | female | 98.31% | 7.7 |
| 322_322a_son | SAMN24905544 | 322 | Adult | 1 | male | 98.59% | 16.6 |
| 322_323_dat | SAMN24905545 | 322 | Adult | 1 | female | 98.49% | 10.8 |
| 322_323a_grndat | SAMN24905546 | 322 | Adult | 2 | female | 99.08% | 16.0 |
| 322_323b_grnson | SAMN24905547 | 322 | Adult | 2 | male | 98.53% | 11.1 |
| 322_323c_grn | SAMN24905548 | 322 | Nymph | 2 | not determined | 98.20% | 9.4 |
| 322_323d_grn | SAMN24905549 | 322 | Nymph | 2 | not determined | 98.40% | 9.6 |
| 338_338_fnd | SAMN24905550 | 338 | Adult | 0 | female | 98.34% | 28.8 |
| 338_338a_son | SAMN24905551 | 338 | Adult | 1 | male | 99.14% | 15.4 |
| 338_339_dat | SAMN24905552 | 338 | Adult | 1 | female | 99.48% | 28.7 |
| 338_339a_grndat | SAMN24905553 | 338 | Adult | 2 | female | 99.48% | 28.3 |
| 338_339b_grnson | SAMN24905554 | 338 | Adult | 2 | male | 98.92% | 13.3 |
| 338_339c_grn | SAMN24905555 | 338 | Nymph | 2 | not determined | 99.11% | 16.4 |
| 338_339d_grn | SAMN24905556 | 338 | Nymph | 2 | not determined | 99.40% | 28.1 |
| 338_339e_grn | SAMN24905557 | 338 | Egg | 2 | not determined | 98.62% | 13.3 |
| 400_400_fnd | SAMN24905558 | 400 | Adult | 0 | female | 99.25% | 22.2 |
| 400_400a_son | SAMN24905559 | 400 | Adult | 1 | male | 99.22% | 25.1 |
| 400_401_dat | SAMN24905560 | 400 | Adult | 1 | female | 99.13% | 28.3 |
| 400_401a_grnson | SAMN24905561 | 400 | Adult | 2 | male | 99.31% | 29.1 |
| 412_412_fnd | SAMN24905562 | 412 | Adult | 0 | female | 99.34% | 22.2 |
| 412_412a_son | SAMN24905563 | 412 | Adult | 1 | male | 99.30% | 25.7 |
| 412_413_dat | SAMN24905564 | 412 | Adult | 1 | female | 99.28% | 19.9 |
| 412_413a_grnson | SAMN24905565 | 412 | Adult | 2 | male | 98.56% | 12.2 |
| 412_413b_grn | SAMN24905566 | 412 | Nymph | 2 | not determined | 98.63% | 13.9 |
| 426_426_fnd | SAMN24905567 | 426 | Adult | 0 | female | 99.08% | 14.4 |
| 426_426a_son | SAMN24905568 | 426 | Adult | 1 | male | 98.73% | 10.5 |
| 426_427_dat | SAMN24905569 | 426 | Adult | 1 | female | 99.43% | 28.8 |
| 426_427a_grndat | SAMN24905570 | 426 | Adult | 2 | female | 98.82% | 27.7 |
| 426_427b_grnson | SAMN24905571 | 426 | Adult | 2 | male | 98.89% | 16.1 |
| 426_427c_grn | SAMN24905572 | 426 | Nymph | 2 | not determined | 99.35% | 21.2 |
| 426_427d_grn | SAMN24905573 | 426 | Nymph | 2 | not determined | 99.30% | 28.9 |
| 426_427e_grn | SAMN24905574 | 426 | Nymph | 2 | not determined | 99.20% | 20.6 |
| 426_427f_grn | SAMN24905575 | 426 | Nymph | 2 | not determined | 99.50% | 32.4 |
| 43_43_fnd | SAMN24905439 | 43 | Adult | 0 | female | 98.54% | 8.2 |
| 43_43a_son | SAMN24905440 | 43 | Adult | 1 | male | 98.89% | 13.1 |
| 43_43b_sis | SAMN24905441 | 43 | Adult | 1 | female | 99.21% | 17.3 |
| 43_44_dat | SAMN24905442 | 43 | Adult | 1 | female | 99.44% | 29.8 |
| 43_44a_grndat | SAMN24905443 | 43 | Adult | 2 | female | 99.54% | 29.1 |
| 43_44b_grnson | SAMN24905444 | 43 | Adult | 2 | male | 99.12% | 13.9 |
| 43_44c_grn | SAMN24905445 | 43 | Nymph | 2 | not determined | 98.94% | 10.6 |
| 43_44d_grn | SAMN24905446 | 43 | Nymph | 2 | not determined | 99.37% | 21.1 |
| 458_458_fnd | SAMN24905576 | 458 | Adult | 0 | female | 99.36% | 24.4 |
| 458_458a_son | SAMN24905577 | 458 | Adult | 1 | male | 98.72% | 12.2 |
| 458_459_dat | SAMN24905578 | 458 | Adult | 1 | female | 99.22% | 27.9 |
| 458_459a_grnson | SAMN24905579 | 458 | Adult | 2 | male | 99.35% | 31.3 |
| 458_459b_grn | SAMN24905580 | 458 | Nymph | 2 | not determined | 99.37% | 31.9 |
| 458_459c_grn | SAMN24905581 | 458 | Egg | 2 | not determined | 99.35% | 27.2 |
| 46_46_fnd | SAMN24905447 | 46 | Adult | 0 | female | 99.04% | 12.8 |
| 46_46a_son | SAMN24905448 | 46 | Adult | 1 | male | 98.64% | 19.6 |
| 46_46b_sis | SAMN24905449 | 46 | Nymph | 1 | not determined | 99.24% | 14.8 |
| 46_47_dat | SAMN24905450 | 46 | Adult | 1 | female | 99.53% | 28.6 |
| 46_47a_grn | SAMN24905451 | 46 | Nymph | 2 | not determined | 98.95% | 10.1 |
| 46_47b_grn | SAMN24905452 | 46 | Nymph | 2 | not determined | 99.21% | 17.4 |
| 46_47c_grndat | SAMN24905453 | 46 | Adult | 2 | female | 99.42% | 31.2 |
| 46_47d_grnson | SAMN24905454 | 46 | Adult | 2 | male | 99.19% | 18.9 |
| 46_47e_grn | SAMN24905455 | 46 | Nymph | 2 | not determined | 99.42% | 25.2 |
| 46_47f_grn | SAMN24905456 | 46 | Nymph | 2 | not determined | 99.08% | 14.8 |
| 46_47g_grn | SAMN24905457 | 46 | Nymph | 2 | not determined | 99.33% | 21.3 |
| 476_476_fnd | SAMN24905582 | 476 | Adult | 0 | female | 98.98% | 29.3 |
| 476_476a_son | SAMN24905583 | 476 | Adult | 1 | male | 98.66% | 20.6 |

| | | | | | | | |
|---|---|---|---|---|---|---|---|
| 476_477_dat | SAMN24905584 | 476 | Adult | 1 | female | 98.24% | 15.6 |
| 476_477a_grndat | SAMN24905585 | 476 | Adult | 2 | female | 98.85% | 21.2 |
| 476_477b_grnson | SAMN24905586 | 476 | Adult | 2 | male | 98.87% | 29.6 |
| 476_477c_grndat | SAMN24905587 | 476 | Adult | 2 | female | 98.83% | 23.5 |
| 476_477d_grndat | SAMN24905588 | 476 | Adult | 2 | female | 98.92% | 23.9 |
| 476_477e_grn | SAMN24905589 | 476 | Egg | 2 | not determined | 99.20% | 29.0 |
| 478_478_fnd | SAMN24905590 | 478 | Adult | 0 | female | 99.08% | 32.9 |
| 478_478a_son | SAMN24905591 | 478 | Adult | 1 | male | 98.78% | 24.1 |
| 478_479-1_dat | SAMN24905592 | 478 | Adult | 1 | female | 99.03% | 30.6 |
| 478_479-1a_grnson | SAMN24905593 | 478 | Adult | 2 | male | 98.58% | 25.6 |
| 478_479-1b_grndat | SAMN24905594 | 478 | Adult | 2 | female | 98.27% | 15.5 |
| 478_479-1c_grn | SAMN24905595 | 478 | Nymph | 2 | not determined | 97.97% | 11.3 |
| 478_479-1d_grn | SAMN24905596 | 478 | Nymph | 2 | not determined | 98.72% | 22.1 |
| 478_479-1e_grn | SAMN24905597 | 478 | Egg | 2 | not determined | 97.58% | 9.1 |
| 478_479-2_dat | SAMN24905598 | 478 | Adult | 1 | female | 98.63% | 15.3 |
| 478_479-2a_grndat | SAMN24905599 | 478 | Adult | 2 | female | 98.45% | 9.1 |
| 478_479-2b_grnson | SAMN24905600 | 478 | Adult | 2 | male | 97.96% | 9.3 |
| 478_479-2c_grn | SAMN24905601 | 478 | Nymph | 2 | not determined | 98.40% | 5.6 |
| 478_479-2d_grndat | SAMN24905602 | 478 | Adult | 2 | female | 98.20% | 11.0 |
| 48_48_fnd | SAMN24905458 | 48 | Adult | 0 | female | 99.09% | 11.3 |
| 48_48a_son | SAMN24905459 | 48 | Adult | 1 | male | 99.33% | 16.0 |
| 48_48b_sis | SAMN24905460 | 48 | Nymph | 1 | not determined | 99.06% | 12.1 |
| 48_49_dat | SAMN24905461 | 48 | Adult | 1 | female | 99.47% | 29.6 |
| 48_49a_grn | SAMN24905462 | 48 | Nymph | 2 | not determined | 99.46% | 27.6 |
| 48_49b_grn | SAMN24905463 | 48 | Nymph | 2 | not determined | 99.39% | 23.4 |
| 48_49c_grn | SAMN24905464 | 48 | Egg | 2 | not determined | 98.82% | 9.8 |
| 48_49d_grn | SAMN24905465 | 48 | Egg | 2 | not determined | 98.56% | 5.6 |
| 48_49e_grn | SAMN24905466 | 48 | Nymph | 2 | not determined | 98.95% | 16.9 |
| 48_49f_grn | SAMN24905467 | 48 | Nymph | 2 | not determined | 98.83% | 18.5 |
| 48_49g_grn | SAMN24905468 | 48 | Egg | 2 | not determined | 99.39% | 27.0 |
| 498_498_fnd | SAMN24905603 | 498 | Adult | 0 | female | 97.82% | 9.2 |
| 498_498a_son | SAMN24905604 | 498 | Adult | 1 | male | 97.32% | 11.4 |
| 498_499_dat | SAMN24905605 | 498 | Adult | 1 | female | 98.98% | 37.6 |
| 498_499a_grnson | SAMN24905606 | 498 | Adult | 2 | male | 98.81% | 29.0 |
| 498_499b_grndat | SAMN24905607 | 498 | Adult | 2 | female | 98.02% | 13.5 |
| 498_499c_grndat | SAMN24905608 | 498 | Adult | 2 | female | 98.55% | 26.6 |
| 498_499d_grn | SAMN24905609 | 498 | Nymph | 2 | not determined | 98.12% | 18.2 |
| 498_499e_grn | SAMN24905610 | 498 | Egg | 2 | not determined | 97.91% | 15.7 |
| 502_502_fnd | SAMN24905611 | 502 | Adult | 0 | female | 97.75% | 9.8 |
| 502_502a_son | SAMN24905612 | 502 | Adult | 1 | male | 98.02% | 11.1 |
| 502_503_dat | SAMN24905613 | 502 | Adult | 1 | female | 98.08% | 14.0 |
| 502_503a_grndat | SAMN24905614 | 502 | Adult | 2 | female | 98.04% | 11.2 |
| 502_503b_grnson | SAMN24905615 | 502 | Adult | 2 | male | 98.32% | 12.2 |
| 502_503c_grn | SAMN24905616 | 502 | Nymph | 2 | not determined | 99.05% | 29.5 |
| 502_503d_grn | SAMN24905617 | 502 | Nymph | 2 | not determined | 98.22% | 17.9 |
| 502_503e_grn | SAMN24905618 | 502 | Egg | 2 | not determined | 98.94% | 30.2 |
| 502_503f_grndat | SAMN24905619 | 502 | Adult | 2 | female | 98.70% | 20.8 |
| 520_520_fnd | SAMN24905620 | 520 | Adult | 0 | female | 98.74% | 28.2 |
| 520_520a_son | SAMN24905621 | 520 | Adult | 1 | male | 98.08% | 15.6 |
| 520_521_dat | SAMN24905622 | 520 | Adult | 1 | female | 98.11% | 15.8 |
| 520_521a_grn | SAMN24905623 | 520 | Nymph | 2 | not determined | 98.47% | 18.3 |
| 520_521b_grn | SAMN24905624 | 520 | Egg | 2 | not determined | 98.59% | 20.9 |
| 520_521c_grn | SAMN24905625 | 520 | Egg | 2 | not determined | 98.46% | 13.3 |
| 520_521d_grn | SAMN24905626 | 520 | Egg | 2 | not determined | 99.02% | 34.7 |
| 520_521e_grn | SAMN24905627 | 520 | Egg | 2 | not determined | 98.74% | 29.0 |
| 534_534_fnd | SAMN24905642 | 534 | Adult | 0 | female | 98.98% | 18.2 |
| 534_534a_son | SAMN24905643 | 534 | Adult | 1 | male | 99.23% | 17.1 |
| 534_535_1_sis | SAMN24905644 | 534 | Adult | 1 | female | 98.92% | 15.4 |
| 534_535_2_dat | SAMN24905645 | 534 | Adult | 1 | female | 98.68% | 13.2 |
| 534_535_2a_grndat | SAMN24905646 | 534 | Adult | 2 | female | 99.10% | 16.8 |
| 534_535_2b_grn | SAMN24905647 | 534 | Nymph | 2 | not determined | 99.25% | 18.9 |

**Table S3.** Primers used for the sequencing validation using Sanger-sequencing method

| **Accession** | **Gene name** | **chromosome** | **forward primer** | **Reverse primer** | **length (bp)** |
|---|---|---|---|---|---|
| LOC111243621 | alpha-mannosidase 2x-like | NW_019211455.1 | TGAGATGTCAATGGCAGGCA | AGGAAGCAAATTCGGGGATGT | 404 |
| LOC111252938 | 5-oxoprolinase-like | NW_019211459.1 | TCCGCATATGGACTTGCGTT | ATCCCAATTGCTCTGACCCG | 374 |
| LOC111254119 | probable E3 ubiquitin-protein ligase RNF144A-A | NW_019211460.1 | ACCAGATCGCACCCTTATGC | AGATGCTTTGCTATCCGCCC | 245 |

**SI References**

1. Wright, S. Inbreeding and homozygosis. *Proc. Natl. Acad. Sci. U. S. A.* **19**, 411–420 (1933).
